# Gut protective *Klebsiella* species promotes microbiota recovery and pathobiont clearance while preventing inflammation

**DOI:** 10.1101/2023.11.14.566997

**Authors:** Vitor Cabral, Rita A. Oliveira, Margarida B. Correia, Miguel F. Pedro, Carles Ubeda, Karina B. Xavier

## Abstract

The microbiota inhabiting the mammalian gut serves as a protective barrier against pathogen invasion through a mechanism known as colonization resistance. Antibiotic treatments can inadvertently disturb the gut microbiota, compromising colonization resistance and increasing host’s susceptible to infections. Non-*pneumoniae Klebsiella* spp. members of the gut microbiota play a crucial role in colonization resistance and clearance from the gut of pathogenic *Enterobacteriaceae* following antibiotic-induced perturbations. Specifically, *Klebsiella* strain ARO112 a gut microbiota isolate, can effectively resist and clear *Escherichia coli* colonization after antibiotic-induced dysbiosis.

We assessed the potential of *Klebsiella* sp ARO112 to promote clearance of *Enterobacteriaceae* pathobiont Adherent-Invasive *E. coli* (AIEC) in an Inflammatory Bowel Disease (IBD) mouse model susceptible to inflammatory episodes. In antibiotic-treated IBD-predisposed mice infected with the AIEC, *Klebsiella* sp. ARO112 promoted a faster recovery of gut microbiota members potentially involved in butyrate production and accelerated pathobiont clearance. Functionally, ARO112-driven microbiota recovery promoted higher butyrate levels and prevented intestinal inflammation compared to untreated animals. Conversely, treatment with the well-known probiotic *E. coli* Nissle 1917 enhanced AIEC colonization and inflammation. Furthermore, we assessed the safety of ARO112 as a potential next-generation probiotic; phenotypic comparison of ARO112 against closely related *Enterobacteriaceae* revealed its lower pathogenic potential, including being more recalcitrant to antibiotic resistance acquisition.

Overall, our results showing that *Klebsiella* sp. ARO112 can resolve infections while contributing to the promotion of intestinal health, underscore its potential as a biotherapy agent that can disrupt inflammation-treatment-infection cycles. This potential extends beyond IBD patients, encompassing individuals with other inflammatory-based conditions related to microbiota imbalances.

## Introduction

The gut microbiota plays a crucial role in human health and disease. An abundant and diverse intestinal community of microorganisms performs essential functions, including providing colonization resistance to potential pathogenic bacteria^1–4^. This protective capacity can be significantly compromised by environmental perturbations that alter the population composition, such as changes in diet or medical treatments, like antibiotics^5–9^. These shifts in microbial populational dynamics can be challenging to rectify, leaving patients more susceptible to infections and other medical complications^8,10^.

The pursuit of interventions to restore dysbiotic gut microbiotas has gained momentum. Currently, both basic research and clinical studies encompass a range of approaches, from introducing single protective bacterial species to well-defined bacterial communities or even complete microbiota, as in fecal microbiota transplants (FMT)^11^. Although the United States Food and Drug Administration has recently approved FMT for the successful treatment of recurrent *Clostridioides difficile* infections in humans^12^, the undefined nature of these procedures poses safety risks^13,14^. These risks underscore the need for the identification of defined strategies, which could involve either single- or multi-strain bacterial cocktails. Next-generation probiotic bacteria, defined as bacterial species native to the host that upon ingestion and/or colonization provide benefits, offer promise for restoring microbiota functions^15^. This requires an extensive understanding of the benefits, potential drawbacks, and the organism’s safety profile, all of which are important to its therapeutic importance.

Patients with high susceptibility to infections often undergo extensive antibiotic treatments, rendering them more susceptible to recurrent infections^16,17^. These patients require innovative approaches to complement current treatments and break the cycle of disease recurrence. Inflammatory Bowel Diseases (IBD) are among such diseases where extensive antibiotic treatments can render the patient more susceptible to frequent infections and recurrence of acute disease symptoms^18,19^. IBD are a group of multi-factorial pathologies with a complex etiology, where genetic predisposition, gut microbiota, and environmental conditions play a role^20–23^. Three outcomes emerge as key aspects of disease cycles that afflict IBD patients: gut microbiota imbalances^24^, increased and persistent intestinal inflammation^25^, and susceptibility to intestinal infections^26^. To manage inflammation flares, IBD patients are often prescribed antibiotics or anti-inflammatory medication^27^. However, these treatment options can inadvertently harm the microbiota, often exacerbating the proinflammatory intestinal environment^16^, thus perpetuating the cycle. The consequent loss of natural protection provided by a balanced microbiota often leads to infections by *Enterobacteriaceae* bacteria, like the adherent-invasive *Escherichia coli* (AIEC), which can also trigger inflammatory episodes^28^.

Probiotics are often considered as potential alternative or complementary treatments for intestinal diseases, but their benefits for complex pathologies like IBD have been limited^29^. The probiotic strain *E. coli* Nissle 1917 (EcN) has been proposed as a potential therapy for IBD^30^. However, despite various proposals and even clinical trials^31^, the European Society for Clinical Nutrition and Metabolism, recommends against probiotics for treatment or prevention of Crohn’s Disease^32^, owing to lack of promising results obtained thus far. Therefore, therapeutic probiotics warrant further research, as their efficacy in patients has been inconsistent^33–36^.

In previous work, we identified a commensal intestinal bacterium, *Klebsiella* sp. ARO112 from the *Enterobacteriaceae* family^37^. ARO112 provides partial colonization resistance against an invading *Escherichia coli* K-12 MG1655 strain and *Salmonella enterica* Typhimurium in a streptomycin-induced intestinal dysbiosis murine model^37^. ARO112 can also partially displace the invader *E. coli*, when administered as a treatment, i.e., after colonization by the invading *E. coli*, in the presence but also in the absence of other microbiota members (in ex-germ-free mice). The ability to displace an already established bacterial invader is not a standard feature, even among probiotic strains. Research has shown that commercially available probiotics can partially displace certain pathogenic bacteria but with varying efficacy^38^. Our previous findings have prompted us to test the potential of ARO112 strain as a next-generation probiotic, capable of eliminating colonization by other *Enterobacteriaceae* invading the gut community during dysbiotic events in a disease model.

Here, we assessed the safety of ARO112 strain by evaluating its pathogenic potential for an assembly of pathogenic traits commonly associated with *Enterobacteriaceae* clinical isolates. Our results show that ARO112, compared to other *Enterobacteriaceae,* has low pathogenic potential, including limited efficiency in acquiring or maintaining plasmids carrying antibiotic resistances, and does not induce host inflammation, unlike EcN. Simultaneously, ARO112 displayed a competitive advantage for colonization against different species from the Pseudomonadota phylum (formerly known as Proteobacteria). Its competitive properties and lack of inflammatory potential had a significant impact in an IBD mouse model, where ARO112 exhibited strong potential as a next-generation probiotic, promoting butyrate-producing microbiota recovery from antibiotic-induced dysbiosis, preventing gut inflammation, and reducing infectious AIEC loads, the most common pathobiont in IBD patients^39^.

## Results

### ARO112 encodes fewer predicted pathogenic traits when compared with other *Enterobacteriaceae*

To address the safety of ARO112 to be used as a probiotic, we performed a genome comparison analysis with fifteen *Enterobacteriaceae* strains to assess their pathogenic potential, including five *E. coli* strains, five *Klebsiella pneumoniae* strains, and five non-*pneumoniae Klebsiella* strains, including ARO112 (Supplementary Table 1). These strains were selected to incorporate human (clinical) and murine bacterial isolates, including pathogenic, commensal, and one established probiotic strain, as well as type or laboratory strains for these groups. With this broad strain selection, we expected to involve a wide range of encoded virulence traits and bacteria with different pathogenic potential.

A whole genome-based phylogenetic tree segregates the fifteen strains into three major clades, differentiating among *E. coli*, *K. pneumoniae*, and non-*pneumoniae Klebsiella* strains (Fig. 1a). The non-*pneumoniae Klebsiella* clade shows ARO112 clustering with *Klebsiella grimontii* (Kgr) type strain and close to *K. michiganensis* (Kmi) type strain, and *Klebsiella* MBC022 mouse commensal strain clustering with the *K. oxytoca* DSM5175 (Kox) type strain. This placement of ARO112 closer to Kgr than Kmi was recently shown by others^40,41^, where phylogenetic analysis with new *Klebsiella* isolates placed ARO112 strain close, but not part of, the cluster containing three *K. grimontii* strains^40^. These results showing that ARO112 is phylogenetically closer to *K. grimontii* than *K. michiganensis*, mean that it can no longer be considered a *K. michiganensis*, as initially classified^37^, and raises the hypothesis that ARO112 might be a new species more closely related to *K. grimontii*. In the *E. coli* clade, the probiotic EcN and the pathobiont AIEC (LF82) strains were phylogenetically closer to each other than to both non-pathogenic *E. coli* K-12 (MG1655) and *E. coli* B (EcB) laboratory strains and to the clinical isolate *E. coli* (Ec1898) (Fig. 1a). In the *K. pneumoniae* clade, the three clinical isolates cluster together (MH258, Kp1012, and Kp834), while the other two *K. pneumoniae* strains (Kpn NCTC9633 and ATCC43816) form a distinct cluster.

**Fig. 1.**
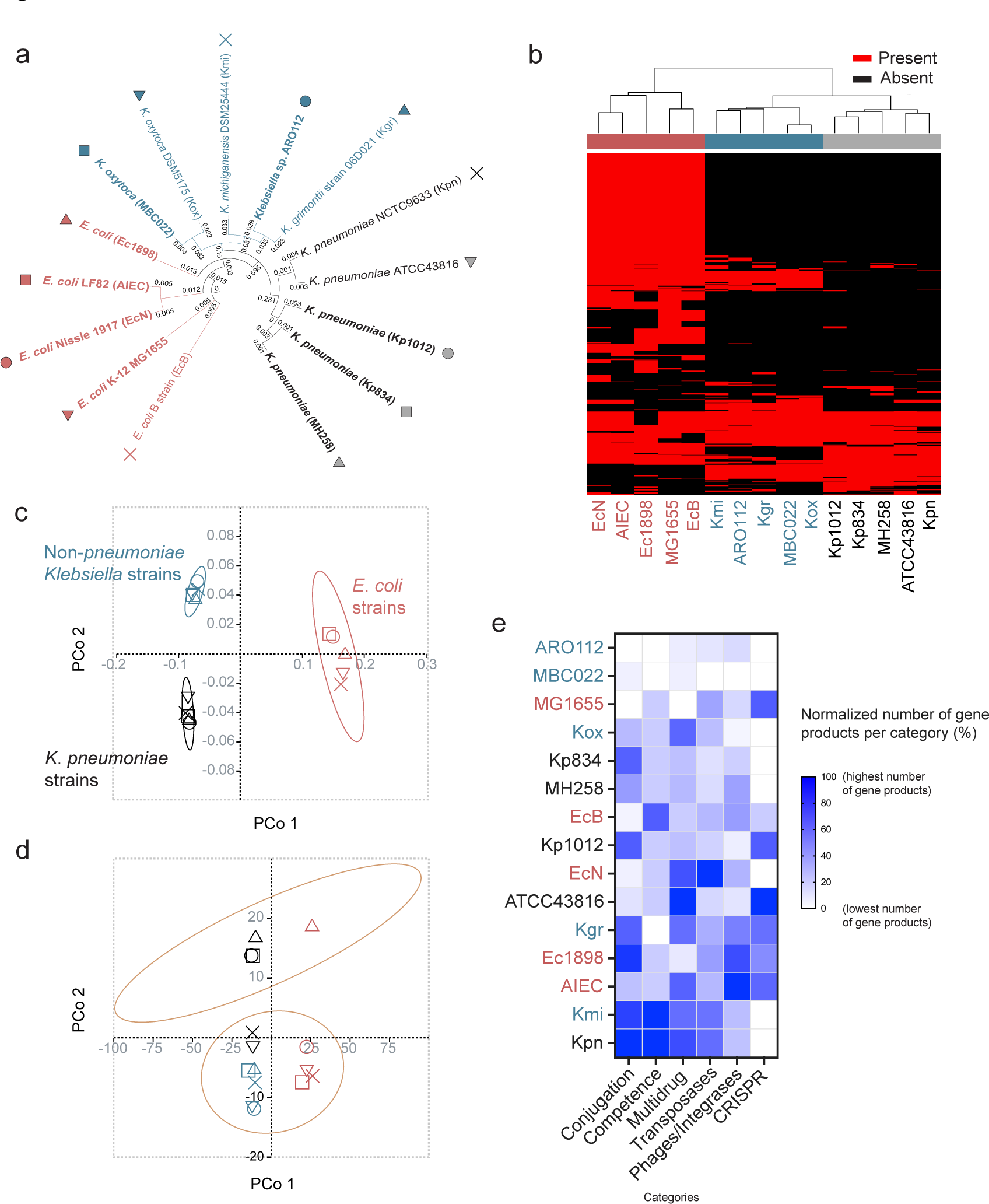
ARO112 genomic comparison with other *Enterobacteriaceae* strains from the *Klebsiella* and *Escherichia* genera for their pathogenic potential. **a,** Phylogenetic tree based on whole-genome analysis of 15 strains (5 *E. coli* strains, 5 *K. pneumoniae* strains, and 5 non-*pneumoniae Klebsiella* strains). Strains in bold were tested experimentally; detailed description of the different the strains in Supplementary Table 1. **b,** Heatmap, **c,** Principal Coordinate Analysis representing presence/absence of pathogenic properties and **d,** Principal Coordinate Analysis of the different strains using the abundance of pathogenic properties. **e,** Normalized quantification of gene products encoding for genes related to clinically-relevant categories (conjugation and conjugal proteins, natural competence, multidrug resistance, transposases, bacteriophages and integrases, CRISPR systems). **c-d,** ellipses represent 90% confidence. **e**, highest number per category is set to 100%, lowest number to 0%, remaining numbers are normalized in between these numbers.

To predict the genome-encoded pathogenic potential of these strains we resorted to 9 databases, reported in Bacterial and Viral Bioinformatics Resource Center^42^, for virulence factors, drug targets and transporters, and antibiotic resistance genes. The presence/absence of the pathogenic properties segregated the three clades represented in the phylogenetic tree (Fig. 1b and 1c). Interestingly, when comparing the ubiquitous gene products and pathogenic properties per clade (shared by the five strains within a clade), there is a higher percentage of gene products that are not ubiquitous to any of the three clades (>75%), compared to the lower percentage of predicted pathogenic traits (<20%) (Supplementary Fig.1).

Upon adding data for the abundance of each pathogenic property (Supplementary Table 4), instead of the binary analysis of presence/absence, the strains were no longer segregated by taxonomy, and instead segregated in two clusters along PCo 2 (Fig. 1d). One cluster included a known multi-drug resistant (MDR) *K. pneumoniae* strain (MH258) and three strains isolated from hospitalized patients. The top 10 pathogenic traits driving the separation of the two clusters include several antimicrobial resistance-associated properties (6/10; Supplementary Table 5), and accordingly these properties were enriched in the cluster including the MDR *K. pneumoniae* strain. ARO112 was in the second cluster together with MBC022, both isolated from non-antibiotic-treated gut microbiomes, and together with known non-MDR strains like MG1655 (Fig. 1d).

To assess the presence of gene products that might have clinical relevance, but not be necessarily included in the available pathogenic properties databases, we analyzed the genomes of each strain for the number of gene products related to the following categories: conjugation, natural competence, multidrug resistance, transposases, phages/integrases, and CRISPR systems. With this analysis, ARO112 consistently shows a lower number of gene products within these categories in comparison with all the other *pneumoniae* and most non-*pneumoniae Klebsiella* strains (Fig. 1e). Importantly, ARO112 was the only *Klebsiella* with no gene products related to conjugation or conjugal proteins, which are important for bacteria to acquire antibiotic resistance genes (Fig. 1e). Moreover, even though the genome analysis displays Kgr as the closest tested relative to ARO112 (Fig. 1a) and the predicted pathogenic traits analysis hints at Kmi as closely related to ARO112 (Fig. 1b), both Kgr and Kmi type strains clearly differ from ARO112 when focusing on gene products for such clinically-relevant categories (Fig. 1e), supporting that ARO112 might have a lower pathogenic potential.

Overall, these results show that the predicted pathogenic properties encoded by the different bacteria correlate with the taxonomy. In contrast, the abundance of these properties segregates by the drug resistance-encoded genes of individual isolates, regardless of taxonomy. ARO112 separated from known MDR strain MH258 and clinical isolates Kp1012, Kp834, and Ec1898, and showed overall less predicted pathogenic properties than the ensemble of the other tested strains.

### ARO112 shows fewer phenotypic traits associated with nosocomial infections

Given that genome predictions do not always directly translate into phenotypes, we decided to perform several in vitro tests for traits that are generally associated with *Enterobacteriaceae* pathogenicity in nosocomial infections and gut colonization (Fig. 2a). For that, we selected a subset of the aforementioned strains, including four *E. coli* strains (non-pathogenic K-12 MG1655 strain, pathobiont AIEC, probiotic EcN, and clinical isolate Ec1898), three *K. pneumoniae* strains (clinical isolates MH258, Kp1012, and Kp834), and two non-*pneumoniae Klebsiella* strains from our strain collection (MBC022, ARO112). We performed these tests under two different laboratory conditions (nutrient rich – LB – and nutrient poor media – M9 salts minimal medium with glucose) to evaluate if these pathogenic traits vary in different environments.

**Fig. 2.**
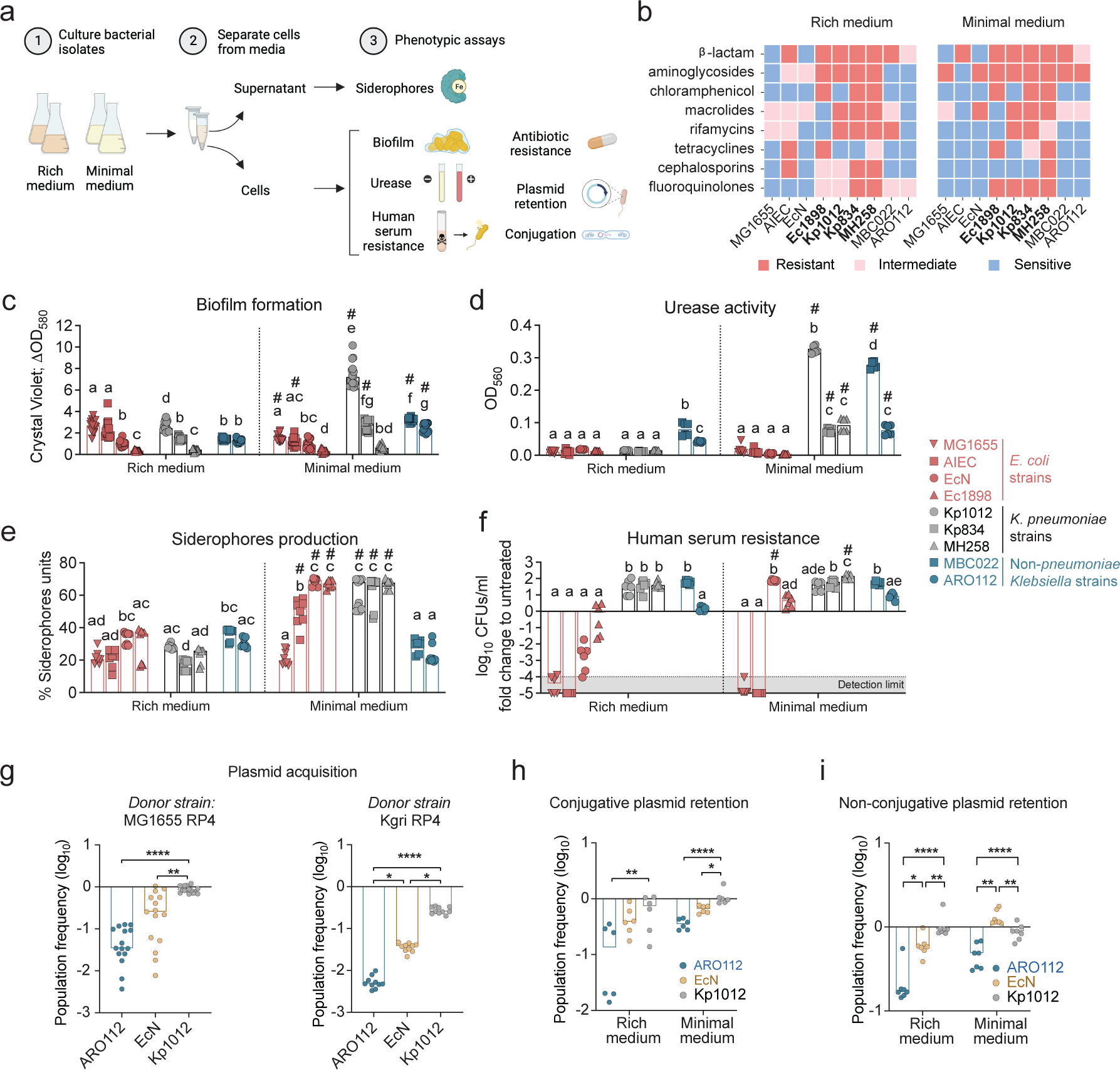
Phenotypical evaluation of ARO112 and other *Enterobacteriaceae* strains for nosocomial-related pathogenic factors important for host invasion and colonization. **a,** Schematic representation of the pipeline used to test bacterial strains for pathogenic traits. A set of 9 strains (4 *E. coli*, 3 *K. pneumoniae*, 2 non-*pneumoniae Klebsiella*) were tested, both in rich and minimal media, for: **b,** antibiotic resistances, MDR *Enterobacteriaceae* strains are represented in bold (resistant to >3 classes of antibiotics); **c,** biofilm formation; **d,** urease activity; **e,** siderophore production; and **f,** capacity to resist killing or inhibition by human serum. Selected strains (ARO112, EcN, and Kp1012) were tested for their capacity to **g**, acquire the conjugative plasmid RP4 from MG1655 donor strain or Kgri donor strain, or **h**, retain the plasmid in rich and minimal media. **i**, The capacity to retain the non-conjugative plasmid pMP7605 in rich and minimal media was also tested. In **c**-**i**, bars represent median values. **b,** includes data from a total of 6 replicates per group, from 2 independent experiments; **c**, from a total of 15 replicates per group, from 3 independent experiments; **d**, from a total of 6 replicates per group, from 2 independent experiments; **e,** more than 8 replicates per group, from 3 independent experiments; **f,** from a total of 6 replicates per group, from 2 independent experiments; **g,** from a total of 15 (from MG1655 RP4) or 10 (From Kgri RP4) replicates per group, from 2-4 independent experiments; **h,** from a total of 6 replicates per group, from 2 independent experiments; **i,** a total of 7 replicates per group, from 2 independent experiments. In **c-g,** statistical analysis was done using Kruskal-Wallis’ test with Dunn’s correction for multiple comparisons. In **e-f,** different letters denote significant differences among strains within the same medium (p<0.05) and # denotes significant differences between different media for the same strain (p<0.05). In **h** and **i**, data were analyzed using Two-way ANOVA with Sidak’s correction for multiple comparisons. * p<0.05; ** p<0.01; **** p<0.0001.

We started by testing antibiotic resistances, as this is the main driver of clinical problems associated with *Enterobacteriaceae*^43^, due to their efficient ability to acquire and share antibiotic resistance genes from other bacteria through mechanisms like horizontal gene transfer (HGT)^44^. Our results showed that ARO112, like MG1655, EcN, and MBC022, is not MDR since it displays resistance to less than three classes of antibiotics (Fig. 2b). The other five strains tested showed resistance to three classes of antibiotics (AIEC in rich medium) or more than three in both media, including, as expected, the known MDR strain MH258 in addition to the clinical isolates from hospitalized patients Ec1898, Kp1012, and Kp834 (Fig. 2b), supporting their clustering together as predicted MDR strains (Fig. 1d). Most of all, these results confirm that ARO112 is not MDR and highlight that bacterial phenotypes are context dependent, with minimal medium, frequently, potentiating resistance phenotypes.

Biofilms are microbial communities enclosed in extracellular matrices and adherent to a substrate, and can contribute to increased recalcitrance to antibiotic treatments^45^. We found that Kp1012 was the strain that produced the most robust biofilm biomass, in minimal medium. In rich medium, most strains resulted in low (EcN, Kp834, MBC022, ARO112) or very low (Ec1898, MH258) biofilm biomass (Fig. 2c), with only MG1655, AIEC, and Kp1012 showing slightly higher biofilm biomass (Fig. 2c). Under minimal medium, ARO112, Kp834, and MBC022 presented some capacity for biofilm formation, but much lower than Kp1012. All *E. coli* strains and MH258 showed lower biofilm biomasses (Fig. 2c).

Urease production is a common trait in *K. pneumonia* and *K. oxytoca* and can be problematic in urinary tract infections, whilst it might also be a good competitive trait for gut colonization^46^. In cultures grown in rich medium, there was no detectable urease activity, except for *K. oxytoca* MBC022 and, to a lesser extent, ARO112, both displaying low levels of urease (Fig. 2d). In cultures grown in minimal medium all *Klebsiella* strains, but not *E. coli* strains, showed significantly higher urease activity, with Kp1012 and MBC022 presenting the highest and Kp834, MH258, and ARO112 the lowest activities, with the latter displaying similar levels to rich medium (Fig. 2d). The absence of urease activity in the *E. coli* strains is expected, since this species is a known non-urease producer, with only some clinical isolates of the pathogenic Enterohemorrhagic *E. coli* reported to have urease activity^47^. Strains proficient in both biofilm formation and urease production, as is the case for Kp1012, can become problematic and often associated with recurrent urinary tract infections^48^.

In rich medium, which is not limited in iron, all strains showed a reduced production of iron chelators, siderophores (Fig. 2e). Interestingly, in iron-limiting conditions, siderophore production was not increased in ARO112, MG1655, nor MBC022, as opposed to the rest of the strains, where a significantly higher siderophore production was observed (Fig. 2e), even though growth yield was unaffected for ARO112, when comparing rich and minimal medium (Extended Data Fig. 1a).

When testing resistance to serum-mediated killing of bacteria grown in rich medium, only the *K. pneumoniae* strains and MBC022 showed significant resistance to and replication ability in the presence of human serum (Fig. 2f), indicating their potential capacity to survive and thrive in the bloodstream of patients. On the contrary, most *E. coli* strains (MG1655, AIEC, EcN) are very susceptible to human serum, while Ec1898 and ARO112 are not killed by nor do they replicate in human serum (Fig. 2f). Interestingly, testing these strains grown in minimal medium resulted in a significantly higher replication for MH258, increasing from 40- (in rich medium) to 140-fold. More surprising, however, was the change detected for EcN where we observed a strong serum resistance and replication capacity when grown in minimal medium (70-fold increase; Fig. 2f). Reports on this phenotype are usually performed using rich media to grow bacteria^49,50^, but here we found that some bacteria, like EcN and MH258, show increased proliferation when grown in minimal medium. Finally, ARO112 shows no evidence of being killed by the human serum, but neither does it replicate to the levels seen for most strains (Fig. 2f). Since this pathogenic trait directly affects the host, as it potentially allows bacteria to proliferate in the bloodstream, with life-threatening risks, ARO112 inability to proliferate is a desirable trait for a potential probiotic.

The evaluation of the pathogenic potential based on phenotypic assays gives us the resolution to the strain level for the pathogenic traits selected. While the genome-based approach showed us that the pathogenic traits predicted to be present or absent in the genome correlate with the taxonomy, the selective testing of clinically-important phenotypes allows us to distinguish distinctly active pathogenic traits within a taxon, such as the heterogeneity within both the *E. coli* and the *K. pneumoniae* clades, but also to pinpoint strains with overall less pathogenic traits associated with nosocomial infections. Notably, ARO112 stands out with low pathogenic traits commonly associated with nosocomial infections.

### ARO112 is resistant to acquire and retain plasmids bearing antibiotic resistance

The lack of resistance found in the commensal non-clinical strains does not exclude the potential for antibiotic resistance acquisition by HGT, which is common among *Enterobacteriaceae* within complex communities, like the gut microbiota^51,52^. Indeed, the acquisition of antibiotic resistance, in particular by HGT events, has been shown to be problematic, with *Enterobacteriaceae* at the forefront of this increasingly concerning medical issue^44^. Therefore, we assessed the capacity of ARO112 to acquire resistances from donor bacteria through HGT, in comparison with the probiotic EcN and the MDR *K. pneumoniae* strain that had typical phenotypes found in causative agents of nosocomial infections, Kp1012. We tested the transfer of a commonly used conjugative plasmid (RP4) from a donor strain (MG1655 or Kgr) into the aforementioned recipient strains. Interestingly, the already MDR strain Kp1012 had the highest acquisition rate, with a median of 83% and 25% of cells acquiring the plasmid from the MG1655 and Kgr donors, respectively (Fig. 2g). EcN presented an intermediate transfer rate, with a median of 25% and 4% acquiring the plasmid and consequent antibiotic resistance from MG1655 and Kgr, respectively (Fig. 2g). In contrast, ARO112 showed strong resistance to plasmid acquisition, with a median of less than 4% of the population acquiring the RP4 plasmid from MG1655 donor strain, and less than 0.5% from Kgr donor strain (Fig. 2g). Next, we tested the capacity to retain the acquired plasmids, after five days of daily passages in rich or minimal media in the absence of antibiotic selection. Remarkably, ARO112 showed the highest plasmid loss, with 87% and 65% of the population losing the plasmid in rich and minimal media, respectively (Fig. 2h). In contrast, the MDR strain Kp1012 lost the plasmid in 27% and 2% of the population, in rich and minimal media, respectively, and EcN lost the plasmid in 61% and 34% of the population in rich and minimal media, respectively (Fig. 2h).

Even plasmids that are non-self-transmissible, can be transferred among bacteria, through conjugation events^53^ in which non-conjugative plasmids take advantage of conjugative plasmids’ machinery. We have also tested the capacity of ARO112, EcN, and Kp1012 to retain a non-conjugative plasmid (pMP7605) carrying an antibiotic resistance received by electroporation. Kp1012 barely lost the plasmid, both in rich (7%) and minimal media (11%; Fig. 2i), and 42% of EcN colonies lost the non-conjugative plasmid in rich medium (Fig. 2i), while no loss was observed in minimal medium (Fig. 2i). Remarkably, ARO112 showed the highest plasmid loss, with 83% and 51% of the population losing the non-conjugative plasmid when grown in rich or minimal media, respectively (Fig. 2i).

Natural plasmids are known to be more stably maintained as they often confer some fitness advantages that drive their persistence (Alonso-del Valle, 2021). We also tested the retention capacity of Kanamycin resistance transferred from one of the MDR clinical isolates predicted to bear natural plasmid(s), Kp834, into ARO112 and EcN (Kp1012 was not tested due to its MDR complicating counterselection). EcN did not lose the resistance, while ARO112 lost the resistance in over 15% (rich medium) and 30% (minimal medium) of the population (Extended Data Fig. 1b).

In summary, even though ARO112 shares with other *Enterobacteriaceae* strains the capacity to acquire plasmids, when compared to the probiotic strain EcN and the *K. pneumoniae* clinical isolate Kp1012, ARO112 is much less efficient in plasmid acquisition and retention, a main source of antimicrobial resistance in *Enterobacteriaceae* and, therefore, presents a reduced risk of becoming a clinical problem.

### Colonization of the murine gut can alter expression of bacterial phenotypic traits

Having observed that the tested phenotypes varied depending on the culture medium used, we questioned which of the tested laboratory media, rich or minimal, was resembling the phenotypical profiles of bacteria in the mammalian gut more closely. Therefore, we tested the phenotypes whose assays do not require bacterial growth, testing directly in bacteria collected from fecal samples of mono-colonized mice (Fig. 3a). We compared the phenotypes of ARO112 with EcN and Kp1012 collected from fecal samples of ex-germ-free mice colonized for five days with each of these bacteria, without selection or growth. Regarding urease activity, Kp1012 presented the same high urease activity detected in minimal medium-grown cultures and EcN showed the expected low urease activity also observed in all laboratory-grown cultures (Fig. 3b and 2d); ARO112 presents a variable phenotype, with half the samples showing low urease activity, similar to the results obtained with cultures grown in the laboratory, while the other half showed increased urease activity (Fig. 3b and 2d). This result, while surprising due to the low levels of urease activity in culture-grown ARO112, indicates that in this strain this phenotype is regulated differently in the gut environment. Interestingly, fecal siderophore measurements showed low values for all three strains, but with a similar tendency observed in minimal medium of lower values for ARO112, when compared with both EcN and Kp1012 (Extended Data Fig. 2a and Fig. 2e).

**Fig. 3.**
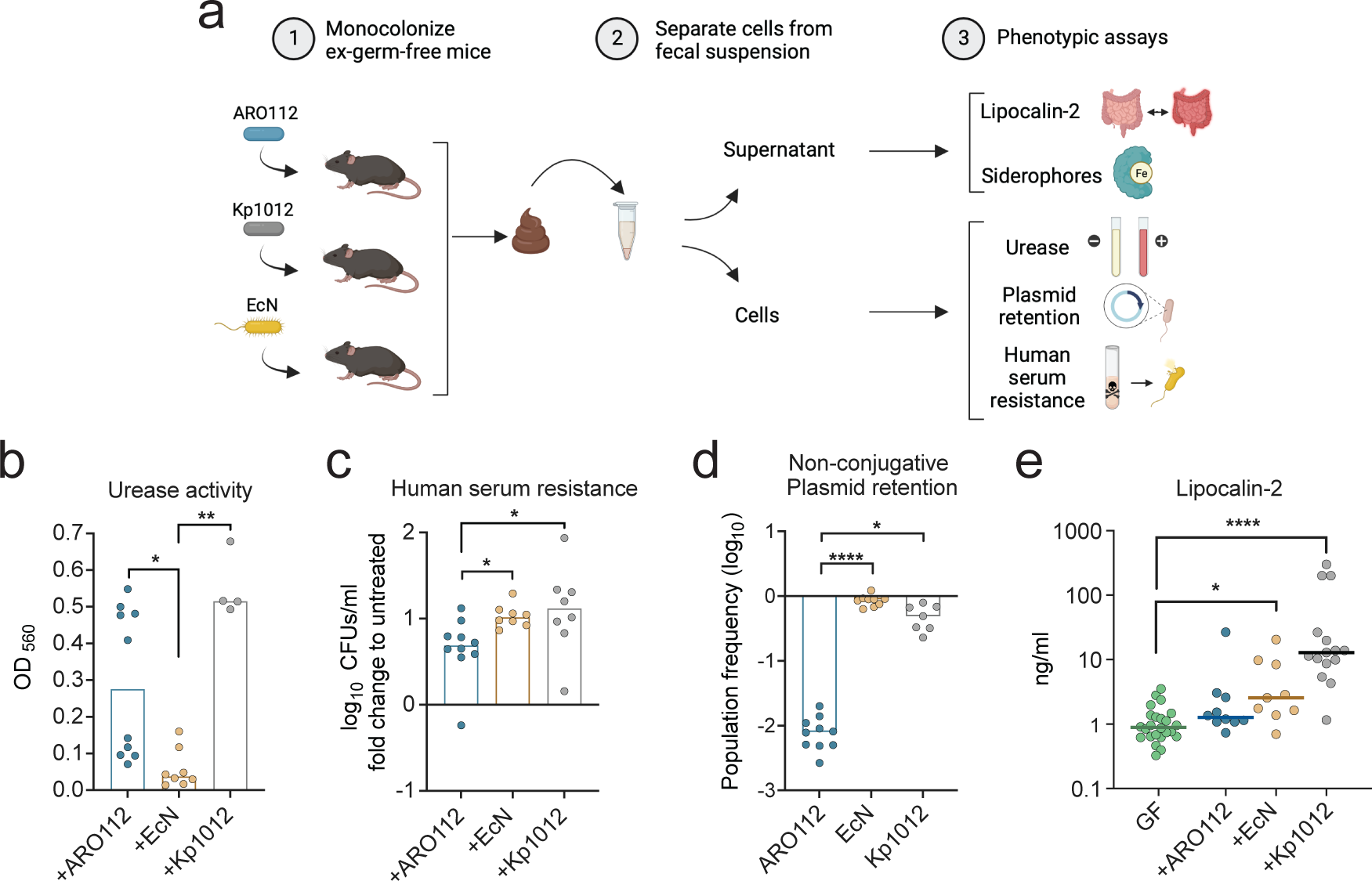
Phenotypic testing of ARO112, EcN, and Kp1012 isolated from fecal samples of mono-colonized mice. **a,** Schematic representation of the pipeline used to test bacterial strains for pathogenic traits after gut colonization. Germ-free (GF) mice were mono-colonized with each of selected strains (ARO112, EcN, Kp1012) for five days, after which bacteria were isolated from the fecal samples and tested for **b,** urease activity and **c,** resistance to human serum. **d,** Bacteria were also evaluated for plasmid retention upon intestinal colonization. **e,** Fecal supernatants were evaluated for Lipocalin-2/NGAL levels, as a marker for intestinal inflammation. In **b**-**e**, data were analyzed using Kruskal-Wallis’ test with Dunn’s correction for multiple comparisons (* p<0.05; ** p<0.01; **** p<0.0001). Urease activity and human serum resistance from gut isolated bacteria were tested in a total of 10 (ARO112), 8 (EcN), or 4-7 (Kp1012) replicates, from at least 2-3 independent experiments. Lcn2 was tested in germ-free animals before colonization (GF, N= 24), and after colonization with ARO112 (N=10), EcN (N=9), or Kp1012 (N=15), from at least 3 independent experiments. Non-conjugative plasmid retention in gut-isolated bacteria was tested in a total of 10 (ARO112), 9 (EcN), or 7 (Kp1012) replicates, from at least 3 independent experiments. In **b**-**e**, bars and lines represent median values.

When testing for resistance to human serum after gut colonization, similar results were obtained with bacteria extracted directly from feces of mono-colonized mice (Fig. 3c) to the ones of laboratory-grown cultures (Fig. 2f). When compared to both Kp1012 and EcN, ARO112 strain is more sensitive to human serum, even after colonizing the mouse gut (Fig. 3c), therefore being a safer strain for clinical settings than the probiotic EcN.

We also checked the non-conjugative plasmid retention in the murine gut by having each strain colonizing germ-free mice carrying the non-conjugative plasmid pMP7605, and quantifying how much of the population retained the plasmid after five days of colonization. EcN lost the plasmid in only 11% of the population, while Kp1012 lost in 51% (Fig. 3d). Remarkably, ARO112 lost the plasmid in more than 99% of cells (Fig. 3d), showing that the recalcitrance of ARO112 to maintaining acquired antibiotic resistances is potentiated in the murine gut.

Next, we tested the host reaction to the colonization by each of the three aforementioned strains, by assessing the fecal levels of Lipocalin-2 (Lcn2), as a marker for gut inflammation^54,55^. After 5 days of mono-colonization, fecal levels of Lcn2 in mice colonized with ARO112 were similar to germ-free mice, providing no evidence for ARO112 inducing intestinal inflammation in mono-colonized mice (Fig. 3e). This contrasted with colonization with EcN and, to an even greater extent, with Kp1012, which significantly induced Lcn2 when compared to the germ-free mice levels (Fig. 3e). Since Lcn2 decreases bacterial growth through competition for iron by siderophore sequestration, we then tested how these three strains were able to maintain growth with increasing levels of Lcn2, which is commonly increased in the inflamed intestinal environment. ARO112, EcN, or Kp1012 were grown in rich or minimal media in the absence or in the presence of increasing concentrations of Lcn2. ARO112 showed an overall higher resistance than EcN to increased levels of Lcn2 (Extended Data Fig. 2b), suggesting a better ability to survive in an inflamed gut environment.

Together, these results indicate that ARO112 could be a safe next-generation probiotic since it has reduced resistance to human serum and low pathogenic potential. Moreover, it has low plasmid retention, as opposed to other *Enterobacteriaceae* strains. Additionally, ARO112’s ability to endure higher levels of Lcn2 with no inflammatory induction, as opposed to EcN, further highlights how this commensal strain is potentially safer and better suited to compete with invading bacteria in intestinal inflammatory environments.

### ARO112 promotes a faster recovery from AIEC gut infection in an IBD mouse model

Having shown that ARO112 has less potentially harmful features than an MDR *K. pneumoniae* and the probiotic EcN, and that in previous work we also see that ARO112 is able to partially displace *E. coli* MG1655 from the gut of mono-colonized and antibiotic-treated mice^37^, here, we decided to test the potential protective role of ARO112 against AIEC. The pathobiont AIEC is particularly problematic under inflammatory conditions, such as those in IBD patients^56^. Therefore, we reasoned that the low inflammatory potential of ARO112 and its capacity to endure Lcn2 could be beneficial in the IBD context.

It was previously shown that a genetic-based murine model for IBD, Nod2^-/-^ mutant mice, is more susceptible to AIEC colonization after antibiotic treatment, similar to IBD patients, as clearance of this pathobiont is slower in the Nod2^-/-^ animals than in WT animals^57^. Additionally, like IBD patients, Nod2^-/-^ animals are more prone to gut inflammation increases^58,59^. Therefore, we started by comparing susceptibility of WT and Nod2^-/-^ mice to colonization by AIEC in our animal facility. All mice were treated by oral gavage with a combination of two non-absorbed antibiotics used in clinical settings on IBD patients^60^, vancomycin and gentamicin, once daily for three consecutive days^57^ (Fig. 4 and Extended Data Fig. 3a). After antibiotic treatment, mice were gavaged with AIEC, and AIEC colonization was followed through selective plating. As observed by others^57^, Nod2^-/-^ mice were more susceptible to AIEC colonization in comparison with WT mice, with ∼50% of the Nod2^-/-^ mice still being colonized with AIEC after 20 days of infection (Fig. 4b, control animals), while all WT mice had cleared the infection in that same timeframe (Extended Data Fig. 3a, control mice). To test the therapeutic potential of ARO112, one day following AIEC infection, Nod2^-/-^ mice were gavaged with either ARO112 strain or the probiotic EcN strain (Fig. 4a). AIEC decolonization was faster in mice treated with ARO112, with 50% of the animals being AIEC-free at 8 days of infection (7 days of probiotic treatment), compared to mice that received no probiotic treatment (+PBS, control mice), in which 14 days were needed for 50% of the mice to resolve the infection, and 7 out of 15 mice still remained infected at the end of treatment (Fig. 4b,c). Surprisingly, treatment with the probiotic EcN did not help decolonization, but actually resulted in a worse outcome, with all but one tested mice remaining infected with high levels of AIEC for the 20 days of the experiment (Fig. 4b,c). We highlight that EcN failed to colonize the one mouse of this group that displaced AIEC infection and therefore, the displacement observed in this animal is not attributed to the EcN treatment (Extended Data Fig. 3b). When assessing the number of mice that resolve the infection in each condition, it is evident that treatment with EcN promotes a continued infection with AIEC, while the absence of probiotic treatment led to the infection being resolved in half of the population (Fig. 4b,c), thus the outcome obtained with the treatment with ARO112, in which 10 out of 12 mice (>83%) became AIEC- free by the end of the experiment, was more efficient than the control (Fig. 4b). Additionally, we observed that ARO112 is itself cleared from the intestinal microbiota throughout the experiment, after the clearance of AIEC, with a median of 10 days post-AIEC inoculation for the ARO112 loads to drop below detection level in 50% of mice (Extended Data Fig. 3b), having disappeared in 9 out of the 12 mice tested (75%) by the end of the experiment (Extended Data Fig. 3b). The protective effect of ARO112 observed in Nod2^-/-^ mice was not seen in WT mice treated with the same antibiotic regimen, as in these animals AIEC decolonization occurs even without treatment (Extended Data Fig. 3a).

**Fig. 4.**
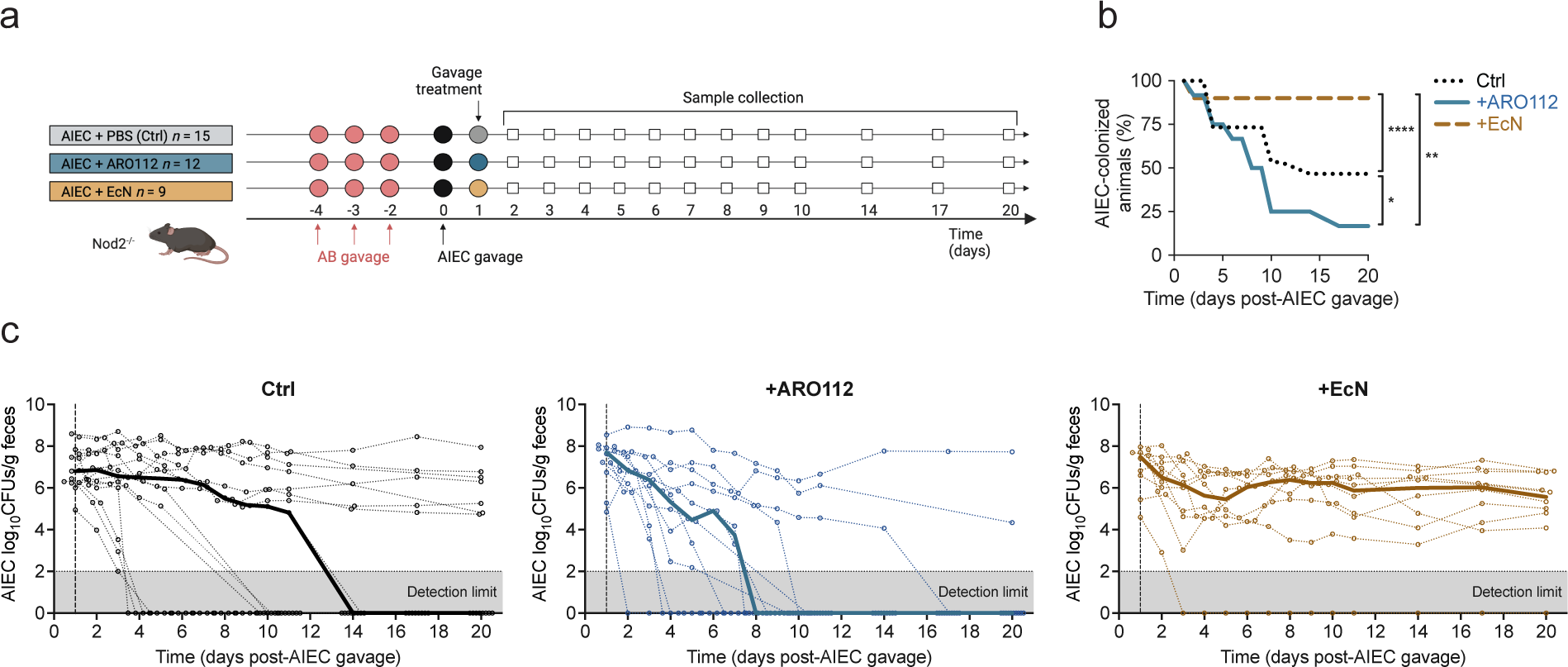
Evaluation of therapeutical potential capacities of ARO112 and EcN against AIEC colonization in an IBD mouse model previously treated with antibiotics. **a,** Experimental setup of IBD mouse model Nod2^-/-^ mice treated with antibiotics, followed by AIEC infection and treated with ARO112, EcN or PBS (Ctrl). **b,** Percentage of mice colonized with AIEC in the different groups throughout the experiment. **c,** AIEC loads during the experiment. Thin lines represent AIEC loads in individual mice over time, while thick lines represent median values for each treatment over time. In **b** and **c**, data were analyzed using Two-way ANOVA with Tukey’s correction for multiple comparisons (* p<0.05; ** p<0.01; **** p<0.0001). AIEC fecal loads and colonization were tested in a total of 15 (Ctrl), 12 (+ARO112), or 10 (+EcN) animals, from 2-3 independent experiments.

We have previously shown that ARO112 could partially displace MG1655 in WT mice with a more severe antibiotic treatment of two weeks of streptomycin in the drinking water^37^, which drastically affects the microbiota composition^61^ and facilitates *Enterobacteriaceae* colonization that compete for the available carbon sources. In streptomycin-treated animals, ARO112 reduced MG1655 colonization largely by a direct interaction between these 2 species through nutrient competition, as the ARO112-dependent inhibition of MG1655 could be observed with laboratory cultures and in germ-free mono-colonized mice^37^. We tested the effect of ARO112 and EcN on AIEC in streptomycin-treated WT mice where, in contrast to what we had observed with MG1655, neither ARO112 nor EcN were able to displace AIEC (Extended Data Fig. 3c). Moreover, in the absence of a microbiome in mono-colonized mice with AIEC, a limited displacement of AIEC was obtained upon co-colonization with ARO112 (∼4 fold; Extended Data Fig. 3d), which was similar to what was observed when co-colonized with other *Enterobacteriaceae* like the MDR clinical isolates Ec1898 and Kp1012 (>4 and ∼6 fold, respectively; Extended Data Fig. 3e,f), but lower than the >10 fold seen previously against MG1655^37^ or against other Pseudomonadota species like *Vibrio cholerae* (>38; Extended Data Fig. 3g). Interestingly, when co-cultured in vitro in static conditions in minimal medium to test for competition for carbon sources, ARO112 also had a minimal effect on AIEC growth as opposed to EcN that seemed to highly reduce AIEC loads (Extended Data Fig. 3h). Together, these results showing that ARO112 weakly inhibits AIEC in vitro, in the absence of other microbiota members (germ-free) or in the presence of low complex microbiota (after streptomycin treatment), indicating that the interaction between these two bacteria is not driven by nutrition competition. Therefore, the mechanisms explaining the displacement of AIEC in the Nod2^-/-^ animals do not seem to rely on a direct ARO112 effect on AIEC colonization and rather indicate the potential involvement of the gut microbiota. Additionally, the above results highlight that ARO112 can have a broad range effect on different Pseudomonadota pathogens, but the mechanism might be different for different competitors.

### ARO112 treatment promotes microbiota recovery, namely potential butyrate-producing bacteria depleted by antibiotic treatment

To further understand the effect of ARO112 on AIEC in the IBD model, we evaluated a possible role of the gut microbiota in this process. We analyzed the microbiota composition of Nod2^-/-^ mice by 16s rRNA gene sequencing before (pre-AB) and after (post-AB) antibiotic treatment, as well as post-AIEC colonization and after 20 days of treatments with PBS, EcN, or ARO112. We observed that mice treated with ARO112 showed a significant recovery in richness and a tendency in recovery of diversity, unlike the long-lasting decrease of both metrics in the mice that received no probiotic treatment or EcN, which persisted as low as just after the antibiotic treatment (Fig. 5a,b). This microbiota recovery promoted by the treatment with ARO112 can help explain both the better displacement of AIEC in this group, but also the displacement of ARO112 itself (Extended Data Fig. 3b), as we expect the recovering microbiota to develop colonization resistance.

**Fig. 5.**
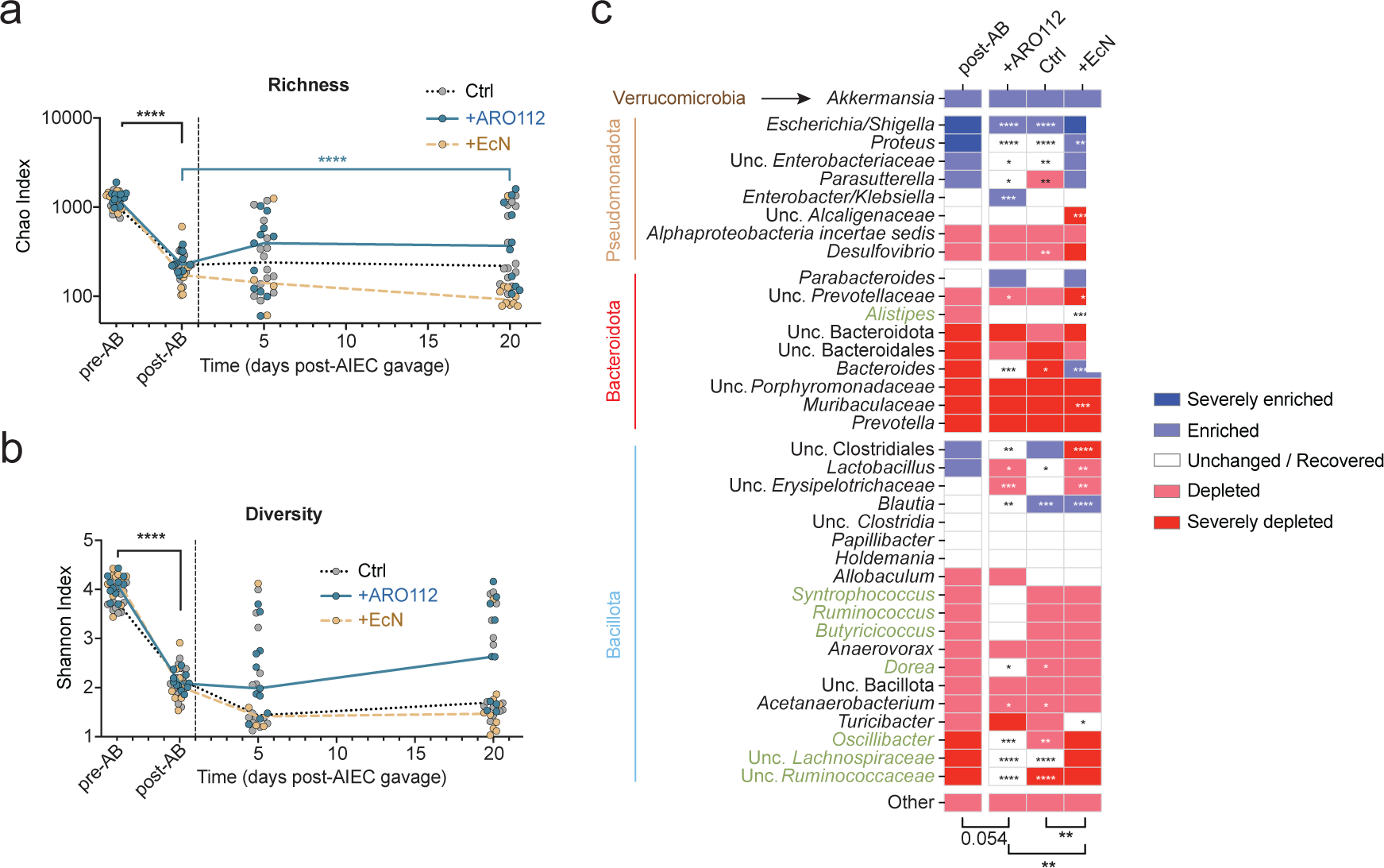
Intestinal microbiota composition resultant from treatment with probiotic strains (ARO112, EcN) or lack thereof (Ctrl). **a,** Richness and **b,** diversity of gut microbiota in fecal samples collected before (pre-AB, day-4), or after (post-AB, day 0) antibiotic treatment and 5 and 20 days after AIEC infections of mice untreated (Ctrl) or treated with ARO112 or EcN. In **a** and **b**, lines represent median values for each treatment over time. **c,** Changes in the relative abundances of most prevalent taxa at day 20 after AIEC infection in comparison with the levels before antibiotics (day-4). In **a**-**c**, data were analyzed using Two-way ANOVA with Dunnett’s correction for multiple comparisons (* p<0.1; ** p<0.01; *** p<0.001; **** p<0.0001). Fecal microbiota composition analyses were performed in a total of 14 (Ctrl), 10 (+ARO112), or 4-10 (+EcN) samples, from 2-4 independent experiments.

We then analyzed which of the most abundant taxa (37 taxa representing an average of 98% abundance of all taxa) were affected by the antibiotic treatment (Fig. 5c). Out of these 37 taxa, 22 (59%) were depleted by the antibiotics, while 7 (19%) were enriched and 8 (22%) remained unchanged by the end of the experiment. The mice from the control group or treated with EcN were able to recover 14% (3 of 22) of taxa depleted by the antibiotic treatment (Fig. 5c). More taxa recovered in the group treated with ARO112 strain, where mice were able to recover 41% (9 of 22) taxa depleted by the antibiotic treatment (Fig. 5c).

A more detailed analysis of the groups of bacteria affected by antibiotic treatment showed that of the 8 taxa of potential butyrate producers^62^ depleted by the antibiotics (highlighted in green in Fig. 5c), 4 recovered upon 19 days post-treatment with ARO112 (*Dorea*, *Lachnospiraceae*, *Oscillibacter* and *Ruminococcaceae*), while only 1 was recovered in the control (*Lachnospiraceae*) and another with EcN treatment (*Alistipes*) (Fig. 5c).

A Principal Coordinate Analysis (PCoA) depicting the Bray-Curtis dissimilarity index with the 37 taxa shows that antibiotic treatment significantly changes the microbiota (pre-AB vs post-AB, Extended Data Fig. 4a). In all three groups the microbiota composition remained different from pre-AB at day 20 of the experiment (Extended Data Fig. 4a), however, treatment with ARO112 resulted in samples positioning closer to the pre-AB levels (Extended Data Fig. 4b). Moreover, a subset of the microbiota appears to be recovered by ARO112 treatment, the potential butyrate producers that are strongly depleted by antibiotic treatment (Fig. 5c). Bray-Curtis dissimilarity indices with the 8 taxa for potential butyrate producers depleted by antibiotics (*Alistipes*, *Syntrophococcus*, *Ruminococcus*, *Butyricicoccus*, *Dorea*, *Oscillibacter*, unclassified *Lachnospiraceae*, unclassified *Ruminococcaceae*) show that, even though both the PBS and ARO112 treatments resulted in microbiomes with a composition significantly closer to the pre-AB levels (Extended Data Fig. 4c), only the population of samples treated with ARO112 is undistinguishable from pre-AB samples, demonstrating an overall recovery of this group of taxa promoted by ARO112 (Extended Data Fig. 4d).

In summary, here we show that in a mouse model susceptible to infection and prone to inflammation, ARO112 promotes recovery of the dysbiotic microbiota by increasing a group of microbiota members potentially involved in butyrate production.

### Probiotic therapy with ARO112 enhances microbiota production of butyrate and prevents intestinal inflammation

As described in the above section, treatment with ARO112 leads to a better recovery of microbiota richness and diversity, including potential butyrate-producing bacteria. Studies have tagged short-chain fatty acids (SCFAs) like butyrate, acetate, and propionate as important metabolites in IBD, with depletion of butyrate-producing gut bacteria by antibiotic treatments being considered a clinical outcome in these patients^16,63^. Moreover, oral administration of butyrate to improve IBD outcomes in patients has been previously proposed^64^. Therefore, our observation that ARO112 treatment promotes potential butyrate producers in the microbiota, possibly leading to an increased and sustained production of butyrate within the microbiota and gut epithelium, instead of the need for continuous exogenous administration, could be very relevant. For that reason, we have assessed SCFA concentrations in fecal samples of mice treated with probiotics or untreated (Fig. 6a). In agreement with our previous results showing the increase of potential butyrate-producing bacteria, 19 days of treatment with ARO112 resulted in a clear increase of fecal levels of butyrate when compared to mice treated with PBS or with EcN (Fig. 6a). An increase in isobutyrate (an isomer of butyrate) and acetate was also observed with the ARO112 treatment in comparison with the control group and EcN-treated mice, although not statistically significant. Propionate levels were significantly higher with ARO112 than with PBS, but similar to the levels in EcN group. On the other hand, succinate showed higher, but not significant, values in mice treated with EcN, when compared to mice treated with PBS or ARO112 probiotic (Fig. 6a).Butyrate has been linked to intestinal health, with anti-inflammatory properties^65^, while succinate has been touted as an inflammation marker^66^ and can accumulate during *Enterobacteriaceae* infections^67^.

**Fig. 6.**
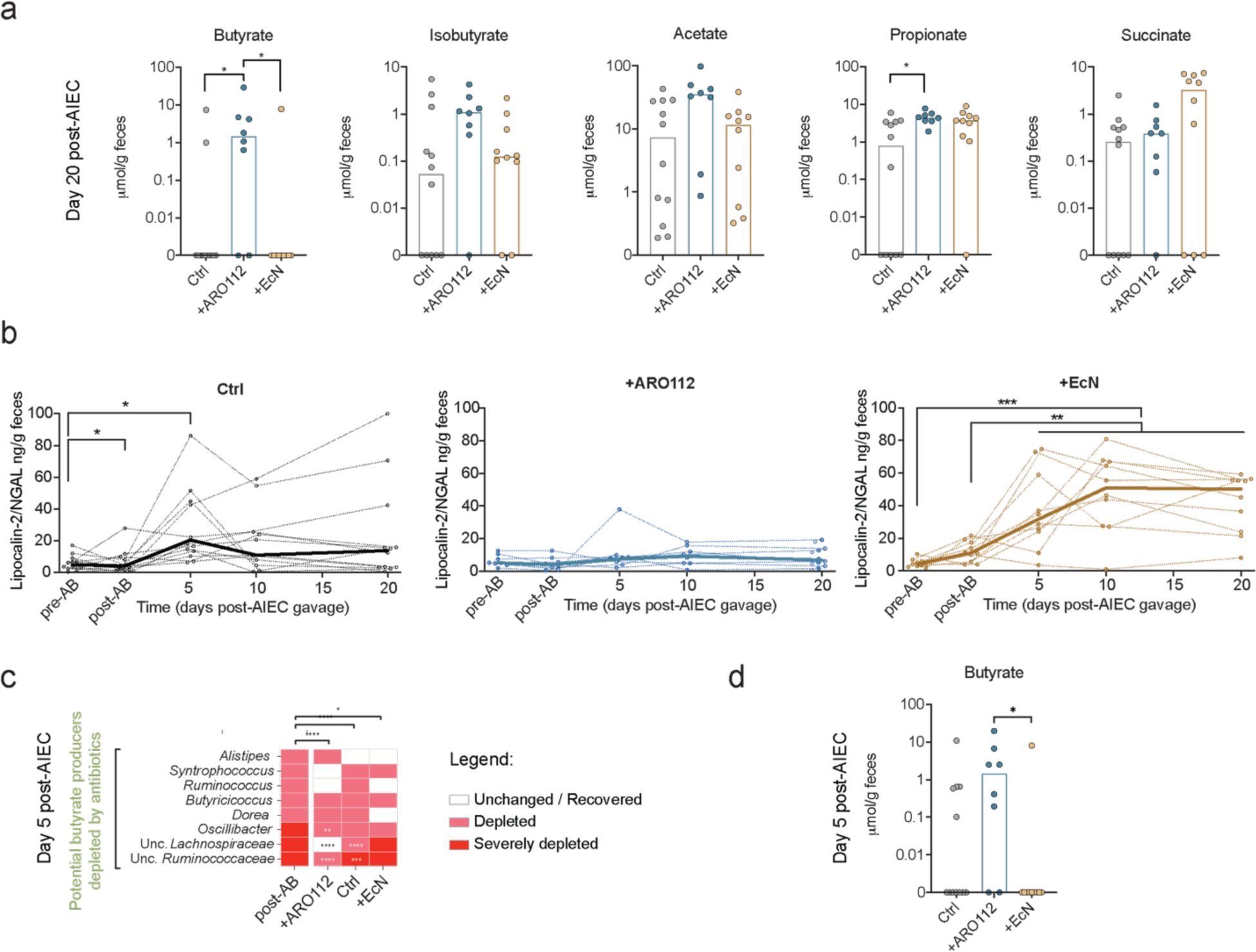
SCFA levels and intestinal inflammation profiles of mice upon different probiotic treatments (ARO112, EcN) or lack thereof (Ctrl). **a,** Metabolomic profile of butyrate, isobutyrate, acetate, propionate, and succinate in fecal samples of IBD mouse model Nod2^-/-^ mice not treated (Ctrl) or treated with probiotics ARO112 (+ARO112) or EcN (+EcN), at the end of the experiment. **b,** Intestinal inflammation was measured via Lcn2 proxy in fecal supernatants, throughout the experiment. Thin lines represent Lcn2 levels in individual mice overtime, while thick lines represent median values for each treatment overtime. **c,** Changes in relative abundances of potential butyrate producers at day 5 after AIEC infection that were depleted by antibiotics, but recovered during experiment. **d,** Metabolomic profile of butyrate in fecal samples of IBD mouse model Nod2^-/-^ mice not treated (Ctrl) or treated with probiotics ARO112 (+ARO112) or EcN (+EcN), at day 5 post-infection (day 4 of treatment). In **a** and **d**, bars represent median values and data were analyzed using Kruskal-Wallis test with Dunn’s correction for multiple comparisons. In **b**, data were analyzed using Two-way ANOVA with Tukey’s correction for multiple comparisons (* p<0.05; ** p<0.01; *** p<0.001; **** p<0.0001). Metabolomics and Lcn2 were tested in 12 (Ctrl), 8 (+ARO112) and 10 (+EcN) samples. In **c**, different colors display log2 fold changes to pre-AB levels of each mouse – severely depleted (<-5), depleted (<-1), unchanged or recovered (−1 to 1), enriched (>1) and severely enriched (>5), and data were analyzed using Two-way ANOVA with Dunnett’s correction for multiple comparisons (* p<0.1; ** p<0.01; *** p<0.001; **** p<0.0001). Fecal microbiota composition analyses were performed in a total of 14 (Ctrl), 10 (+ARO112), or 4- 10 (+EcN) samples, from 2-4 independent experiments.

Next, we used Lcn2 to evaluate the intestinal inflammatory response of the Nod2^-/-^ mice infected with AIEC and treated with PBS, ARO112, or EcN. Interestingly, treatment with EcN not only led to persistent infection by AIEC but also resulted in persistently high Lcn2 levels (Fig. 6b), unlike PBS treatment, in which Lcn2 levels varied between animals, having a significant but mostly transient increase at day 5 of infection (Fig. 6b). More importantly, mice treated with ARO112 showed persistently low levels of Lcn2, never significantly varying from the levels prior to antibiotic treatment (Fig. 6b).

The reduced Lcn2 levels and increased butyrate observed here upon treatment with ARO112 are in accordance with previous reports on the anti-inflammatory properties of butyrate^65,68^. However, because the preventive effect on inflammation promoted by ARO112 treatment was seen as early as 5 days post-infection, for it to be due to the anti-inflammatory properties of butyrate would require recovery of butyrate producers early in the treatment. Therefore, we analyzed the state of recovery of potential butyrate producers depleted by antibiotics at day 5 of infection (4 days of treatment). Results show that ARO112 treatment resulted in a significant recovery of three potential butyrate producers, one of which, unclassified *Lachnospiraceae*, already recovered to levels seen before antibiotic treatment (Fig. 6c). To evaluate if this recovery would translate in higher levels of intestinal butyrate, we also measured butyrate levels at day 5, noticing that butyrate levels in the gut of mice treated with ARO112 already showed higher levels than the other two groups. We reason that this early recovery of butyrate production promoted by ARO112 can explain the prevention of inflammatory episodes at that early stage of treatment.

These results show that ARO112 treatment promotes recovery of a subset of microbiota, namely butyrate producers depleted by antibiotics, which through increased production of butyrate prevent inflammation, leading to pathogen clearance and better recovery. Moreover, the possible pro-inflammatory potential of EcN also highlights ARO112 as a safer probiotic in both non-inflammatory and inflammatory host states. The ensemble of these results, namely the displacement of AIEC, the prevention of inflammation, and the recovery of the dysbiotic microbiota and butyrate levels, all obtained upon treatment with ARO112, show a three-pronged therapy targeting the hallmarks of IBD (infection, inflammation, and dysbiosis).

## Discussion

In this study, we show that the microbiota member of the non-*pneumoniae Klebsiella* complex, *Klebsiella* sp. ARO112 strain, is able to displace infections by the pathobiont AIEC in an IBD mouse model. Remarkably, this displacement was not merely driven by a competitive interaction between ARO112 and AIEC. Instead, ARO112 therapy after antibiotics treatment facilitated microbiota recovery, promoting specific members, like *Lachnospiraceae*, and enhancing butyrate production. This recovery had functional consequences and benefits beyond just AIEC displacement, also preventing increased intestinal inflammation during infection.

Other recent studies have added encouraging evidence that non-pneumoniae *Klebsiella* microbiota species and strains contribute significantly to protection against *Enterobacteriaceae* infections, demonstrated in animal models and supported by human studies^37,40,41^. One study pinpointed and validated *K. oxytoca* strains present in children’s microbiota with protection against invading *K. pneumoniae*^40^. There, the authors showed that nutrition competition played an important role in protection against *K. pneumoniae*, similarly to what we had already shown in a previous publication where ARO112 protected against *E. coli* and *Salmonella* Typhimurium through nutrient competition^37^. A more recent publication studying human samples identified a series of bacterial sequences matching non-*pneumoniae Klebsiella* species (related to *K. michiganensis*, *K. oxytoca*, and ARO112) associated with protection against intestinal pathogens with high risk of bacteremia^41^. These three studies support the protective role of non-*pneumoniae Klebsiella* species against *Enterobacteriaceae* invasion, highlighting the role of nutrient competition in this interaction. In contrast, the results from our present study show that direct effects of ARO112 against *E. coli* is strain-dependent, since ARO112 shows less capacity to displace AIEC and MRE strains, than it had shown to displace MG1655^37^ or *V. cholerae* in the absence of other microbiota, which also shows that the ability of ARO112 to displace pathogens is not restricted to *Enterobacteriaceae*. This indicates that nutrition competition and inhibitory mechanisms potentially involved in the interactions between these pairs of bacteria are not relevant under the conditions tested, supporting our proposal that ARO112 displacement of AIEC in the IBD mouse model is mediated by its promotion of microbiota recovery. Displacement based on direct interactions, although likely to be useful in many situations, are case specific as they depend on genomic profiles of the protective-pathogen pair^69,70^. On the other hand, the ability of ARO112 to promote displacement through microbiota recovery has broader applications, potentially being effective against a wider range of pathogenic strains as the exact genomic composition of the pathogen is less relevant.

Despite the encouraging support for the role of these protective non-*pneumoniae Klebsiella* in colonization resistance, the fact that these bacteria are also *Enterobacteriaceae* raises concerns for clinical applications. *Enterobacteriaceae* pathogens are the major cause of Hospital infections in Europe^71^. Remarkably, genome analysis of ARO112 in comparison with other *Enterobacteriaceae* revealed that ARO112 has less predicted pathogenic traits. The low pathogenic potential based on predicted functions was supported by our phenotypic tests for traits associated with nosocomial infections. Our results of phenotypic tests for pathogenic traits show that some phenotypes are context-dependent, highlighting the importance of testing cells cultured in different media and, if possible, cells extracted from colonization experiments, before asserting the safety of a specific strain for therapeutic purposes. Notably, ARO112 demonstrated to have a lower pathogenic potential in comparison with the patient isolate Kp1012, and similar or lower than the probiotic EcN, which expresses several traits that are commonly associated with pathogenesis, as previously reported^35,36^.

In Hospitals, the biggest concern related to *Enterobacteriaceae* infections is the high prevalence of MDR^72^, which result in critical clinical situations due to a lack of available alternative treatments. We showed that ARO112 not only is not MDR but also showed a low capacity for acquiring and retaining resistant plasmids carrying antibiotic resistance. Interestingly, genome analysis of ARO112 revealed no evidence for genes predicted to be involved in conjugation, which is a major mechanism for plasmid acquisition in *Enterobacteriaceae*^73^.

The link between ARO112’s ability to displace AIEC and to promote microbiota recovery is likely tied to its impact on potential butyrate producers, namely on the recovery of members of the *Lachnospiraceae* family and the increase of butyrate levels, which were observed as early as 5 days after infection in ARO112- treated animals. Butyrate is known to be important for gut health due to its role in promoting epithelial metabolism and immune regulation and having anti-inflammatory properties^65,74,75^. Butyrate has also been shown to provide colonization resistance to *Salmonella* infections^76,77^ and MDR *Enterobacteriaceae*^78,79^ by direct inhibition. Similarly, studies have shown the importance of *Lachnospiraceae* in colonization resistance against multiple pathogens vancomycin-resistant enterococci (VRE)^80^, *Clostridium difficile*^81^, *Salmonella* Typhimurium^70^, and *Listeria monocytogenes*^82^. In a previous study, we observed that the presence of microbiota members from *Lactobacillus* genus could restrict MRE colonization by promoting the expansion of Clostridiales, with an increase in butyrate levels observed in both mice and antibiotic-treated leukemia patients^79^. However, in that study, even though addition of *Lactobacillus* could decrease MRE gut invasion by two orders of magnitude, complete clearance was never obtained. Additionally, cocktails containing *Lactobacillus* are often used as probiotics but can delay natural microbiota compositional recovery in healthy individuals^33^. In contrast, the effect obtained by ARO112 treatment in the IBD mouse model resulted in clearance of the pathobiont AIEC with a success rate of more than 80%, alongside an accelerated microbiota recovery.

Other studies also support the notion that many mechanisms of colonization resistance require positive interactions among unrelated microbiota members to promote colonization resistance and pathogen clearance^79,80^. Here, we describe an innovative ARO112 therapeutic effect that goes beyond infection clearance, showing that in an IBD model, ARO112 promotes the recovery of native butyrate producers depleted by antibiotic treatment while preventing inflammatory flares in the process. Therefore, the therapeutic effect of ARO112 is multifactorial, affecting all three main hallmarks of intestinal inflammatory disorders: susceptibility to infection, dysbiosis, and gut inflammation.

Currently available probiotics therapies often show low efficacy largely due to common caveats: 1) unable to colonize long enough to actuate, 2) cause delay in natural microbiota recovery and 3) contribute to the increase of antibiotic resistance transfer^33,34,38,83^. However, ARO112 strain does not display any of these caveats in the IBD murine model studied here. ARO112 not only does not delay microbiota recovery, but actually promotes recovery while also having the advantage of being cleared from the gut in the process. This is in striking contrast with what happened with the probiotic EcN, which delayed microbiota recovery, prolonged AIEC colonization and persisted in the gut at high loads, resulting in increased levels of gut inflammation.

Having a safe protective microbiota member that can be used to promote a faster recovery of an imbalanced microbiota and displace pathogens or pathobionts, while preventing inflammatory flares in a disease context is of great therapeutic importance. This safe and effective approach opens new avenues for probiotic research and therapeutic development, offering potential solutions for a range of other clinically-relevant disorders. Further exploration of ARO112’s protective mechanisms under different contexts will advance our understanding and enable the identification of additional safe therapeutic microbiota members or consortia for clinical use.

## Acknowledgements

We thank the members of the Bacterial Signalling Laboratory at Instituto Gulbenkian Ciência and Tanja Dapa, and João Xavier for critically reading the manuscript. In particular, we thank Joana Amaro for the help processing samples for genomic sequencing and metabolomics. We also thank Instituto Gulbenkian Ciência’s Animal Facility, in particular Joana Bom, and the Genomics Facility, namely João Sobral, for the help provided during this project’s execution. The authors would like to thank the Microbiology Service from the Hospital La Fé in Valência (Spain) for isolating and providing the strains Ec1898, Kp834, and Kp1012. I would like to thank Gabriel Nuñez for providing the AIEC strain (LF82), and Roberto Balbontín-Soria for providing the plasmid pMP7605. We would also like to acknowledge Athanasios Typas for the suggestion to test conjugation from a clinical isolate.

This work was funded by Fundação para a Ciência e Tecnologia (FCT-Portugal) projects [PTDC/BIA- MIC/6990/2020] and [SFRH/BPD/116806/2016], as well as by Municipality of Oeiras Proof-of-Concept InnOValley 2022 [IOVPoC-2022-21] and a European Commission grant [MSCA-IF-2018-843183]. Additionally, this project was also funded by InfectERA-ERANET- through FCT-Portugal grant (InfectERA/0004/2015) and Acciones complementarias grant [PCIN-2015-094] from the Spanish Ministerio de Economía y Competitividad and the 7th Research framework program from EU, and a grant from the Spanish MICINN [PID2020- 120292RB-I00], granted to KBX and CU, respectively. NMR data were acquired at CERMAX, ITQB-NOVA, Oeiras, Portugal with equipment funded by FCT, project AAC 01/SAICT/2016.

## Author contributions

V.C. and R.A.O conceptualized and designed the research studies with advice and support from K.B.X and C.U.. R.A.O. executed in vitro experiments with support from V.C.. V.C. executed in vivo experiments with AIEC and Kp1012, in SPF wildtype, SPF Nod2^-/-^, and GF wildtype models, with support from R.A.O.. M.B.C. performed in vivo experiments with *V. cholerae* and Ec1898 with support from R.A.O.. M.F.P. performed the NMR. R.A.O. analyzed microbiome sequencing data and V.C. performed the post-analysis. C.U. provided the whole-genome sequencing data for human isolates Kp1012, Kp834, Ec1898. V.C. and R.A.O. analyzed and interpreted all the data, under the supervision of K.B.X.. V.C. and R.A.O. wrote the first draft of the manuscript with valuable input from K.B.X. and C.U.. All authors critically read and approved the manuscript.

## Material and Methods

### Bacterial strains, plasmids and culture conditions

See Supplementary Table 1 for all species, strains, and plasmids. Unless otherwise specified bacteria were cultured in Lysogenic Broth (LB, here refereed as rich medium) or M9 minimal medium (47.7mM NaHPO4-7H2O, 22mM KH2PO4, 8.6mM NaCl, 18.7mM NH4Cl, 2mM MgSO4, 0.1mM CaCl2, 1mM thiamine)^84^ with 0.5% glucose at 37°C with aeration (shaking at 240 rpm or in static cultures, as specified).

### Whole genome sequencing

*Klebsiella oxytoca* MBC022 was isolated by plating on LB agar the feces of a WT specific pathogen-free (SPF) C57BL/6J mice bred in the animal house facility at Instituto Gulbenkian Ciência. DNA isolation was performed using a previously described protocol^85^ from cells in liquid LB at 37 °C with agitation. DNA library and whole genome sequence was obtained at the IGC Genomics facility. Paired-end sequencing of each sample was performed using an Illumina MiSeq Benchtop Sequencer (Illumina), which produced datasets of 250 bp read pairs.

Ec1898, Kp1012, and Kp834 were isolated from leukemia patients from Hospital La Fe in Valencia (Spain). Genome sequence was obtained as described above in the FISABIO Sequencing facility.

All sequences have been deposited in GenBank.

### Method for phylogenetic tree

Phylogenetic tree was obtained using the whole genomes of strains uploaded to the PATRIC BV-BRC browser software (v3.29.20). Parameters used included 100 single-copy genes, using Multiple Alignment using Fast Fourier (MAFFT) Transform alignment program and RAxML Fast Bootstrapping branch support method (v8.2.11).

### Genome-encoded predicted pathogenic properties and gene products

Genomes of all strains were uploaded to the PATRIC BV-BRC^42^ browser software (v3.29.20) and analyzed. For each genome, a table containing all predicted pathogenic properties from 9 available databases: 3 for Virulence Factors (Victors, PATRIC_VF, VFDB), 3 for Drug Targets and Transporters (DrugBank, TCDB, TTD), and 3 for antibiotic resistances (PATRIC, CARD, NDARO). The ensemble of these data was downloaded and two merged tables were compiled with the information for all the strains: one displaying the presence or absence of each pathogenic property in each strain, and another including the abundance of each pathogenic property in each strain. Principal Coordinates analyses were performed using either presence/absence of pathogenic properties (Bray-Curtis dissimilarity index for binary/categorical data; one-way PERMANOVA with Bonferroni correction, 999 permutations) or abundance of said pathogenic properties (Euclidean distances for continuous data; one-way PERMANOVA with Bonferroni correction, 999 permutations).

Genome-encoded gene products relating to clinically-relevant categories (conjugation and conjugal proteins, natural competence, multidrug resistance, transposases, bacteriophages and integrases, CRISPR systems) number of hits were obtained (Supplementary Table 3) and normalized in graphpad Prism software: in each category the highest number was listed as 100%, the lowest as 0%, and the other numbers distributed according to the percentage within this interval.

### Antibiotic resistance profile

Cultures were grown in either rich or minimal medium for 24h at 37°C with shaking. Cultures were washed once in sterile PBS and adjusted to a final OD600 of 0.05 in 150μl/well in the appropriate medium with different concentrations of antibiotics (as specified). These cultures were then grown for 24h in a 96-well plate in a plate shaker, after which OD600 measurements were performed in a plate reader Multiskan Sky. Strains were considered resistant if in the presence of antibiotic could reach at least 80% of the growth obtained in the absence of antibiotic. If growth in the presence of antibiotic was more than 40% and less than 80% of the control growth, the strain was considered intermediate for resistance, and sensitive if grown less than 40% of control growth. Resistance or intermediate resistance to one antibiotic of any class is considered resistance or intermediate resistance to that class of antibiotics. Sensitivity to a class of antibiotics means that bacteria were sensitive to all antibiotics tested of that class. A strain was considered multidrug resistant (MDR) if it was non-susceptible (sensitive or intermediate) to at least 1 antimicrobial agent in 3 or more antimicrobial classes, in both tested media, as defined before^17^.

| Class | Antibiotic | Concentration<br>(µg/ml) |
| --- | --- | --- |
| β-lactam | ampicillin | 100 |
|  | carbenicillin | 50 |
| Aminoglycosides | streptomycin | 100 |
|  | kanamycin | 50 |
|  | gentamicin | 15 |
|  | neomycin | 200 |
|  | apramycin | 50 |
|  | spectinomycin | 50 |
| Chloramphenicol | chloramphenicol | 25 |
| Macrolides | erythromycin | 125 |
| Rifamycins | rifampicin | 20 |
| Tetracyclines | tetracycline | 10 |
| Fluoroquinolones | ciprofloxacin | 0.5 |
|  | trimethoprim | 4 |
| Naphthyridone | nalidixic acid | 32 |
| Cephalosporins | cefotaxime | 8 |

### Biofilm formation

Biofilm formation assays were performed by adapting a previously published method (Reisner, 2006; O’Toole, 2011). In summary, bacterial cultures were grown for 24h in either rich or minimal media at 37°C with shacking. The following day, each culture was washed once in sterile PBS, and final OD600 adjusted to 0.05 in the appropriate medium and 200 μl of culture were dispensed per well in 48-well plates. Plates were centrifuged for 5 min at 4000 rpm at room temperature, after which plates were incubated at 37°C for 90 min for adhesion. After adhesion, supernatants were carefully removed and discarded, wells were washed once with 200 μl of sterile PBS, and 500 μl of sterile fresh appropriate medium were carefully dispensed per well. Plates were incubated at 37°C for 24h, after which attached biofilms (biomass) was quantified with Crystal Violet (CV). For that supernatants were removed and discarded and well bottoms were carefully washed with 500 μl of sterile PBS. Each well was carefully stained with 0.1% CV solution and plates were incubated at room temperature for 20 min protected from light. After incubation, each well was washed with PBS and plates were incubated open and inverted in a paper towel for 30 min protected from light, to dry. Each well was then de-stained with 200μl of a 33% glacial acetic acid solution and incubated for 15 min at room temperature protected from light. Supernatants were removed from each well into a new plate and read for OD580 in a Multiskan Sky plate reader (peak for CV staining), to quantify biofilm biomass.

### Urease activity

Cultures were grown in either rich or minimal medium for 24h at 37°C with shaking. Cultures were washed once in sterile PBS and adjusted to an OD600 of 1, after which each culture was diluted 20x in urea medium (3g/L L-tryptophan, 5 g/L NaCl, 1g/L H2KO4P, 1g/L H2K2O4P, 20g/L urea, 0,012g/L phenol red) and incubated for 24h at 37°C with shaking. Urea degradation leads to the medium color to change from yellow/orange to pink/red. 100 μl of each culture were then transferred into a 96-wellplate and measured at OD560 in a Multiskan Sky plate reader to quantify urease activity. For urease activity measurement of bacteria extracted from fecal samples of monocolonized mice, cell suspensions of 2.5×10^7^ cells/ml were used, instead of the grown culture.

### Siderophore production

Cultures were grown in either rich or minimal medium for 24h at 37°C with shaking. Cultures or fecal samples diluted in PBS were centrifuged to recover supernatants. Culture and fecal supernatants were filtered in 0.22 μm filters. Cell-free supernatants (100 μl) were mixed 1:1 (v:v) with Cas-PIPES solution (100μl) (Alexander & Zuberer, 1991; Das & Barooah, 2018) in a 96-well plate and incubated at room temperature for 20 min, after which OD630 was measured in Multiskan Sky plate reader to calculate percentage of siderophore production, using sterile media or PBS as reference.

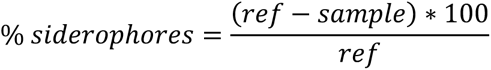

### Resistance to pooled human serum

Bacterial susceptibility to human serum was tested as described (Podschun, 1993) with modifications. Bacterial strains were grown in either rich or minimal medium for 24h at 37°C with shaking, or extracted directly from fecal pellets. Cultures were washed once in sterile PBS and adjusted to 2×10^7^ cells/ml, after which 25 μl of cell suspension in PBS were added to 75 μl of pooled human serum solution (P30-2401, PAN-Biotech). Cultures were plated in LB agar plates for CFUs at 0h and after 3h of static incubation at 37°C, to assess killing, resistance, or proliferation of bacterial cells in the presence of human serum.

### Lipocalin-2 in vitro growth

Cultures were grown in either rich or minimal medium for 24h at 37°C with shaking. Cultures were washed once in sterile PBS and adjusted to a final OD600 of 0.05 in the appropriate medium with different concentrations of Lipocalin-2 protein. Cultures were grown for 24h in a 96-well plate in a plate reader Biotek Synergy H1 with regular OD600 measurements. Final OD600 obtained were plotted against the concentration of Lipocalin-2 protein used.

### Plasmid conjugation

Cultures were grown in rich medium for 24h at 37°C with shaking. Cultures were washed once in sterile PBS. Donor (MG1655 RP4, Kgri RP4) and recipient (ARO112, EcN, Kp1012) strains were thoroughly mixed at a 1:1 ratio (total of 10^8^ cells), centrifuged for 30 sec at 14,000 rpm and resuspended in 20 μl of PBS. Two drops of 10 μl each were placed on an agar plate, air-dried, and incubated at 37°C for 24h. After incubation, both drops were removed with a sterile loop and resuspended in 1 ml of sterile PBS. Serial dilutions were performed and plated to count CFUs for recipient strain with (selective plating with plasmid-borne antibiotic resistance) and without (non-selective or selective plating for recipient’s antibiotic resistance plating) tested plasmid. Conjugative plasmid (RP4) harbors resistances to tetracycline (10 μg/ml) and kanamycin (50 μg/ml).

### Plasmid retention

Strains harboring tested plasmids (RP4 acquired through conjugation from MG1655 RP4 donor strain; pMP7605 acquired through electroporation; Kanamycin resistance acquired through conjugation from clinical isolate Kp824) were grown in either rich or minimal medium with antibiotics (for plasmid-borne antibiotic resistances; kanamycin (50 μg/ml) and tetracycline (10 μg/ml) for RP4; gentamicin (30 μg/ml) for pMP7605; kanamycin (50 μg/ml) for transfer from clinical isolate) for 24h at 37°C with shaking. Cultures were washed once in sterile PBS and adjusted to a final OD600 of 0.05 in 150 μl in the appropriate medium without antibiotics, and serial dilutions were plated in selective and non-selective LB agar plates, to assess CFUs with (plasmid retention) and without (plasmid loss) antibiotic resistance. Cultures were incubated at 37°C with shaking for 24h. Every 24h, cultures were diluted 1:100 in fresh appropriate medium and re-incubated. After 5 days of passages, serial dilutions of cultures were plated as mentioned before.

### Animal experimentation

All of the mouse experiments performed at Instituto Gulbenkian Ciência (IGC) were approved by the Institutional Ethics Committee and the Portuguese National Entity (Direção Geral de Alimentação e Veterinária; 015190), which complies with European Directive 86/609/EEC of the European Council.

C57BL/6J male mice, either WT or Nod2^-/-^, were used at 6–8 weeks of age and were randomly assigned to experimental and control groups. C57BL/6J mice were used for all experiments. None of the animal experiments were performed blinded. Sample size was chosen according to institutional directives and in accordance with the guiding principles underpinning humane use of animals in research. No statistical analyses were performed to predetermine the sample sizes. All of the experiments were performed at least twice, except when stated otherwise.

### Bacterial isolation from fecal samples

WT C57BL/6J germ-free mice were orally gavaged with 100 μL containing ∼10^8^ CFUs of ARO112, EcN, or Kp1012 strains bearing the non-conjugative plasmid pMP7605. Five days later, fecal pellets were collected for live gut bacteria extraction, as previously described^51^ (Ronda, 2019). Briefly, pellets were weighed and 500 μL of sterile PBS were added before mechanical disruption with a motorized pellet pestle, after which another 500 μL of sterile PBS were added. Samples were subjected to four iterations of vortex mixing for 15 sec, centrifugation at 1,000 rpm for 30 sec at room temperature and recovery of 750 μL of the debris-free cell-containing supernatants into a new tube, replacing that volume with sterile PBS before the next iteration. After recovery of 3 ml per sample, isolated cells were centrifuged at 4,000 g for 5 min at room temperature, discarding the supernatant and resuspending the cells fractions in 1 ml of sterile PBS supplemented with glycerol and cysteine at final concentrations of 20% and 0.1%, respectively. Samples were frozen and stored at −80°C, to measure phenotypes (urease activity, human serum resistance, plasmid retention) as described above. Fecal supernatants were used to assess fecal Lipocalin-2 levels and siderophores.

### Nod2^-/-^ and WT SPF mice experiments

C57BL/6J Nod2^-/-^ and WT mice bred under specific pathogen-free conditions in the animal house facility at the Instituto Gulbenkian Ciência were kept individually in sterile ISOcages (Tecniplast) at our specific opportunistic pathogen-free facility. Mice were orally gavaged with gentamicin (3 mg/kg/d) and vancomycin (40 mg/kg/d) once daily for three consecutive days (Drouet, 2012). Two days after the last gavage with antibiotics, mice were orally gavaged with 100μL containing ∼10^8^ CFUs of AIEC strain (LF82-strepR), and the following day were orally gavaged either with 100 μL of PBS (control), or 100 μL containing ∼10^8^ CFUs of ARO112 or EcN strains. Fecal samples were collected at the timepoints indicated in the corresponding graph and plated in selective medium to assess colonization levels of AIEC. Additionally, fecal samples were also used to measure levels of Lcn2 (Lipocalin-2/NGAL ELISA) and to determine the intestinal microbiota composition and levels of SCFA.

C57BL/6J WT mice bred under SPF conditions in the animal house facility at the Instituto Gulbenkian Ciência were kept individually in ventilated cages with high-efficiency particulate air filters in our animal facility. Streptomycin (5 g/l) was maintained ad libitum in the drinking water for 15 days, being replaced every 3 days, after which mice were kept in non-supplemented water. Four days after halting streptomycin treatment, AIEC strain (LF82-strepR) was orally gavaged (∼10^8^ CFUs/mouse in 100 μL). Three days after that, either ARO112 strain or EcN strain were orally gavaged (∼10^8^ CFUs/mouse in 100 μL). Fecal samples were collected at the timepoints indicated in the corresponding graph and plated in selective medium to assess colonization levels of AIEC.

### WT gnotobiotic mice colonization clearance experiments

C57BL/6J germ-free mice were bred and raised in IGC’s Gnotobiology Unit in axenic isolators (La Calhene/ORM) and were later transferred into sterile ISOcages (Tecniplast). Animals were individually housed and orally gavaged with 100 μL containing ∼10^8^ CFUs of AIEC (LF82-strepR), Kp1012, *V. cholerae*, or Ec1898 strains. The following day, mice were gavaged with 100 μL containing ∼10^8^ CFUs of either ARO112 or EcN strains, or 100µl of PBS. Fecal samples were collected at the timepoints indicated in the corresponding graphs and plated in selective media to assess colonization levels of AIEC, Kp1012, *V. cholerae*, or EC1898 strains.

### 16S rRNA sequencing

All fecal samples from SPF mice experiments stored at −80 °C were processed for DNA extraction as previously described^61^. In brief, DNA was extracted using a combination of the QIAamp Fast DNA Stool Mini Kit (Qiagen) according to the manufacturer’s instructions, mechanical disruption using a motorized pellet pestle and elution to a final volume of 100 μL in ATE buffer. For each sample, the 16S rRNA gene was amplified using the 515F/806R (V4 regions) primer pairs recommended by the Earth Microbiome Project under the following PCR cycling conditions: 94 °C for 3 min; 35 cycles of 94°C for 60s, 50°C for 60s and 72°C for 105s; and extension at 72°C for 10min^86,87^. After library preparation, 2 × 250 bp sequencing was performed at the IGC Genomics Unit using an Illumina MiSeq Benchtop Sequencer.

### Fecal microbiota analysis

Mothur v.1.32.1 was used to process sequences as previously described^61^, with some modifications. Sequences were converted to FASTA format. Sequences that were shorter than 220 bp containing homopolymers of longer than 8 bp or undetermined bases with no exact match with the forward and reverse primers, and barcodes that did not complement each other or that did not align with the appropriate 16S rRNA variable region, were not included in the analysis. A quality score above 30 (range, 0 to 40, where 0 represents an ambiguous base) was used to process sequences, which were trimmed using a sliding-window technique over a 50 bp window. Sequences were trimmed from the 3ʹ end until the quality score criterion was met, and were merged after that. Between 20,000 and 50,000 sequences were obtained per sample. 16S rRNA gene sequences were aligned using SILVA template reference alignment^88^. ChimeraSlayer^89^ was used to remove potential chimeric sequences. Sequences with distance-based similarity of greater than 97% were joined into the same OTU using the average-neighbor algorithm. All of the samples were rarefied to the same number of sequences (10,000) for diversity analyses. Samples below 10,000 reads were removed from the analysis. A Bayesian classifier algorithm with a 60% bootstrap cut-off was used for each sequence^90^; sequences were assigned to the genus level where possible or otherwise to the closest genus-level classification. All OTUs were used, and taxa plotted are the result of the merge of OTUs with the same classification, being collapsed to 37 different taxa, with lower represented OTUs/taxa being summed and plotted as “other”. Principal Coordinate Analyses were performed using the 37 taxa or a subset of 8 taxa of potential butyrate producers depleted by antibiotics (*Alistipes*, *Syntrophococcus*, *Ruminococcus*, *Butyricicoccus*, *Dorea*, *Oscillibacter*, unclassified *Ruminococcacea*, unclassified *Lachnospiraceae*), representing the Bray-Curtis dissimilarity index (one-way PERMANOVA with Bonferroni correction and 999 permutations).

### Lipocalin-2/NGAL enzyme-linked immunoassay (ELISA)

Fecal samples were weighted and resuspended in 1 ml sterile PBS and mechanically homogenized with a motorized pellet pestle. Samples were centrifuged at 4°C for 15 min at 18,000 xg. Supernatants were recovered and filtered in 0.22 μm filters. ELISA for Lipocalin-2/NGAL (Human Lipocalin-2/NGAL DuoSet ELISA) were performed according to manufacture instructions, except for the volumes used, which were adapted for 384-well plates, instead of 96-well plates. The following volumes were used instead: reagent diluent (40μl/well), sample and standards (20 μl/well), Detection Antibody (20 μl/well), Streptavidin-HRP (20 μl/well), Substrate solution (20 μl/well), Stop solution (20 μl/well).

### SCFA quantification

1H-NMR (proton nuclear magnetic resonance) analysis was performed to determine the abundance of SCFA in the fecal samples collected from Nod2^-/-^ mice treated with ARO112 and EcN, or untreated (control), as previously described^91^. Fecal samples (1 to 3 fecal pellets) were diluted in 1mL of PBS and mechanically homogenized. To remove large debris, the samples were pelleted by centrifugation at 18000 g for 15 min at 4°C. Supernatants was collected and filtered through a 0.22 mm filter (Milipore), followed by another filtration step with 3 KDa filters (Vivaspin 500) using centrifugation at 15000 xg and 4°C for 3 h (or until 150 μL of filtrate was obtained). Filtered samples were stored at −80°C until spectra were acquired. For spectrum acquisition, samples were thawed at room temperature and 150 μL of sample mixed with 60 μL of 350 mM phosphate buffer (pH 7.09 with 2% NaN3, 10 μL of a 0.05% (w/v) 3-(Trimethylsilyl)propionic-2,2,3,3-d4 (TSP- d4, Sigma-Aldrich) solution, and 380 μL of deuterated water - D2O) (for a total volume of 600 μL). This mixture was transferred to a 5 mm glass NMR tube. All solutions were prepared with D2O. Samples were homogenized by inversion and the spectra were acquired after pH measurement. Acquisitions were performed on a Bruker NEO 500MHz instrument equipped with QXI H-C/N/P 5 mm probe-head with z-gradients. 1H-NMR spectra were acquired using 1D NOESY pulse sequence with pre-saturation (noesypr1d) under the following conditions: 90 degrees pulse for excitation mixing time 100 ms, acquisition time 4 s, and relaxation delay 1 s. All spectra were acquired with 200 scans at 25°C, with 48,000 data points and 6002 Hz (12 ppm) spectral width (Chenomx acquisition parameters). The recorded 1H-NMR spectra were phase corrected using Bruker TopSpin 4.0.7 and spectra were then analyzed using Chenomx NMR Suite 8.1. Compounds were identified by manually fitting reference peaks to spectra in database Chenomx 500 MHz Version 10. Quantification was based on internal standard peak integration (TSP-d4).

### Graphical representation and Statistical analyses

Phylogenetic tree (Fig. 1a) was done using iTOL (itol.embl.de). Cluster heatmap (Fig. 1b) was done using SRPlot (https://www.bioinformatics.com.cn), using bidirectional clustering, complete cluster method and Euclidean distances. All statistical graphs and statistical analyses presented were done using Graphpad Prism 8. Principal Coordinate Analyses were obtained with Past4.02 software (https://folk.universitetetioslo.no/ohammer/past) as explained above.

## Figures

**Extended Data Fig. 1.**
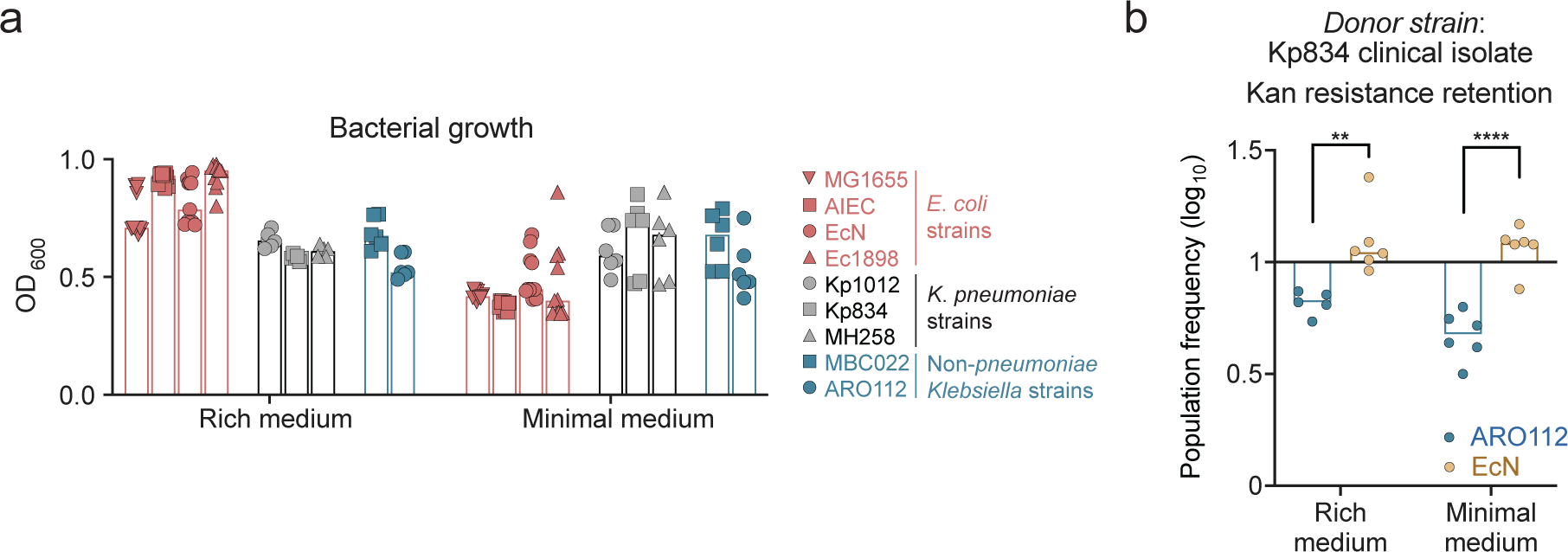
Comparison and correlations of phenotypically tested virulence traits. **a,** Bacterial growth in rich and minimal media. **b,** ARO112 and EcN were tested for their capacity to retain a natural antibiotic-resistant plasmid acquired by conjugation from the clinical isolate Kp834. In **a** and **b**, bars represent median values. In **b**, data were analyzed using Two-way ANOVA test with Sidak’s correction for multiple comparisons (**p<0.01; **** p<0.0001). Growth was tested in 6-11 replicates, from 2 independent experiments. Natural conjugative plasmid retention was tested in a total of 5 replicates per group, from 2 independent experiments.

**Extended Data Fig. 2.**
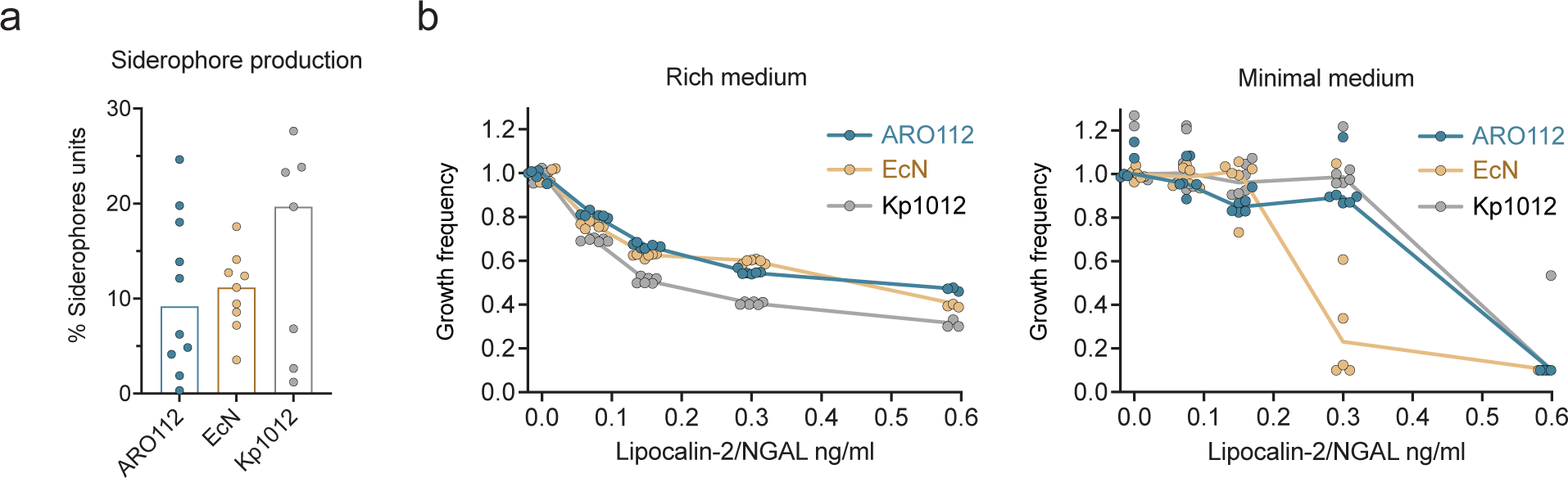
Siderophore production in mono-colonized mice and resistance to Lcn2. **a,** Siderophores were measured in fecal supernatants collected from mice mono-colonized with each of selected strains (ARO112, EcN, Kp1012) for 5 days. Bars represent median values. **b**, Selected strains (ARO112, EcN, Kp1012) were tested for their capacity to grow in the presence of increased concentrations of Lcn2 in rich or minimal media. In **a**, siderophore production in fecal samples was tested in a total of 10 (ARO112), 9 (EcN), or 7 (Kp1012) samples, from at least 3 independent experiments. In **b**, Lcn2 resistance was tested in a total of 6 replicates, from 2 independent experiments, and data were analyzed using Two-way ANOVA with Tukey’s correction for multiple comparisons (* p<0.05; *** p<0.001; **** p<0.0001).

**Extended Data Fig. 3.**
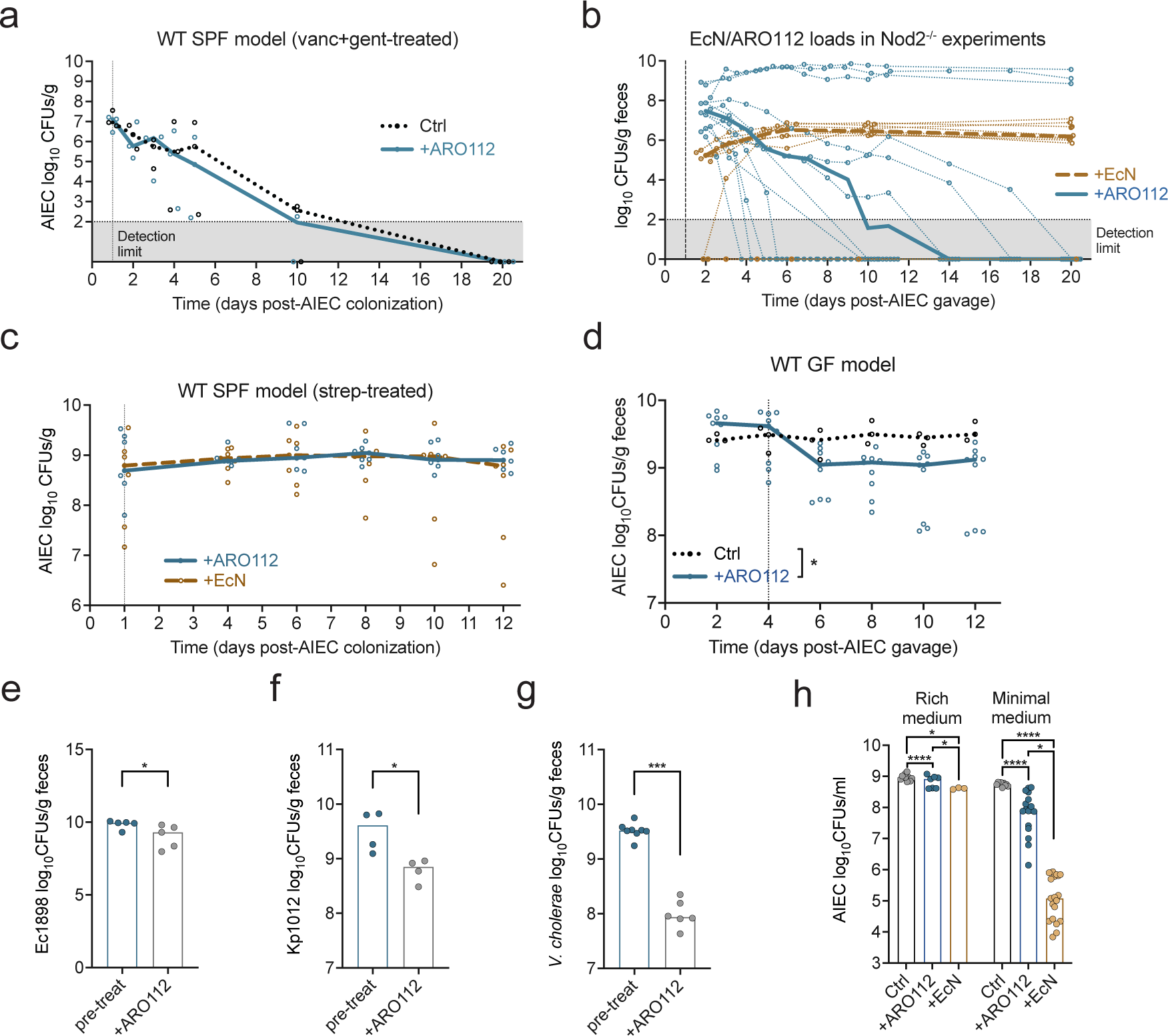
Probiotic colonization and evaluation of protective capacities of ARO112 and EcN against different invading bacteria in different mouse models. **a,** Loads of AIEC in fecal samples of WT mice treated with vancomycin and gentamicin and treated with PBS (Ctrl) or ARO112 probiotic (+ARO112). b, data from experiment shown in Fig. 4, where **b,** shows the loads of ARO112 or EcN in fecal samples. **c,** Loads of AIEC in fecal samples of WT mice treated with streptomycin and treated with ARO112 (+ARO112) or EcN (+EcN) probiotics. **d,** Loads of AIEC in germ-free WT mice treated with PBS (Ctrl) or ARO112 probiotic (+ARO112). Loads of **e,** Ec1898, **f,** Kp1012, and **g,** *V. cholerae* after 4 days of mono-colonization (pre-treat) or after 8 days of treatment with ARO112 (+ARO112), in germ-free WT mice. **h,** Loads of AIEC after 24h of growth without or with ARO112 or EcN, in rich or minimal media. AIEC loads in WT SPF models were tested in 3 replicates per group (**a**) or 7 replicates per group (**c**), from 1 or 2 independent experiments, respectively. Probiotic loads were tested in 12 (+ARO112) or 10 (+EcN) replicates from 2-3 independent experiments (**b**). AIEC loads in WT GF model were tested in 3 (Ctrl) and 9 (+ARO112) replicates in 1-2 independent experiments, and data were analyzed using two-way ANOVA with Sidak’s multiple comparison correction (* p<0.05) (**d**). Ec1898 loads were assessed in 5 replicates from 2 independent experiments (**e**). Kp1012 loads were assessed in 8 replicates from 2 independent experiments (**f**). *V. cholerae* loads were assessed from 6 replicates from 2 independent experiments (**g**). In vitro growths were tested in 10 (Ctrl), 7 (+ARO112), or 3 (+EcN) replicates in minimal medium, and 9 (Ctrl), 19 (+ARO112), or 19 (+EcN) replicates in rich medium, from 1-4 independent experiments (**h**), and two-way ANOVA was tested with Tukey’s multiple comparison correction (* p<0.05; **** p<0.0001). In f-h, data were analyzed using Mann-Whitney tests (* p<0.05; *** p<0.001). In b, thin lines represent ARO112 and EcN loads in individual mice over time, while in a, b, d, e, thick lines and bars represent median values.

**Extended Data Fig. 4.**
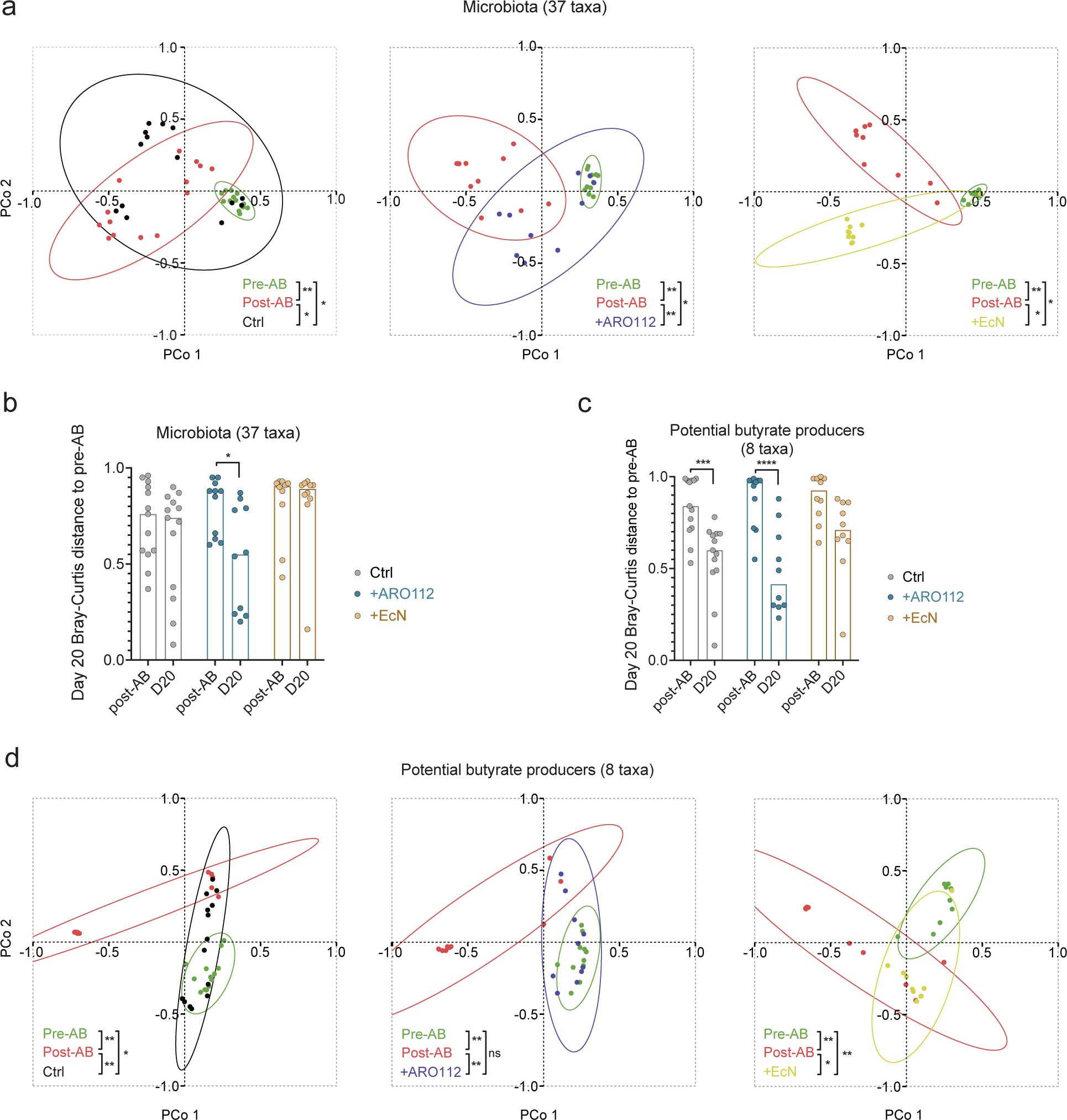
Comparison between treatments of different microbiota-related parameters in experiment in Nod2^-/-^ mice after 19 days of treatment. **a,** Principal Coordinate Analysis of Bray-Curtis dissimilarity index displaying clustering of samples before antibiotic treatment (pre-AB), after antibiotic treatment (post-AB), and after treatment (day 20 of experiment) with PBS (Ctrl; 14 samples) or probiotics (+ARO112 or +EcN; 10 samples per group), using microbiota composition (37 taxa). **b,** Analysis of the Bray-Curtis dissimilarity distances, obtained comparing the samples of each mouse pre-AB, to their position post-AB or post-treatment (day 20). **c,** Analysis of the Bray-Curtis dissimilarity distances, obtained comparing the samples of each mouse pre-AB, to their position post-AB or post-treatment (day 20). **d,** Principal Coordinate Analysis of Bray-Curtis dissimilarity index displaying clustering of samples before antibiotic treatment (pre-AB), after antibiotic treatment (post-AB), and after treatment (day 5 of experiment) with PBS (Ctrl; 14 samples) or probiotics (+ARO112 or +EcN; 10 samples per group), for the 8 taxa of potential butyrate producers depleted by antibiotics (*Alistipes*, *Syntrophococcus*, *Ruminococcus*, *Butyricicoccus*, *Dorea*, *Oscillibacter*, unclassified *Ruminococcacea*, unclassified *Lachnospiraceae*). Ellipses in **a** and **d** represent 90% confidence intervals. One-way PERMANOVA with 999 permutations were performed with Bonferroni corrections (**a** and **d**) and two-way ANOVA were tested with Sidak’s correction for multiple comparisons (**b** and **c**) (* p<0.05; ** p<0.01; *** p<0.001; **** p<0.0001).

**Extended Data Fig. 5.**
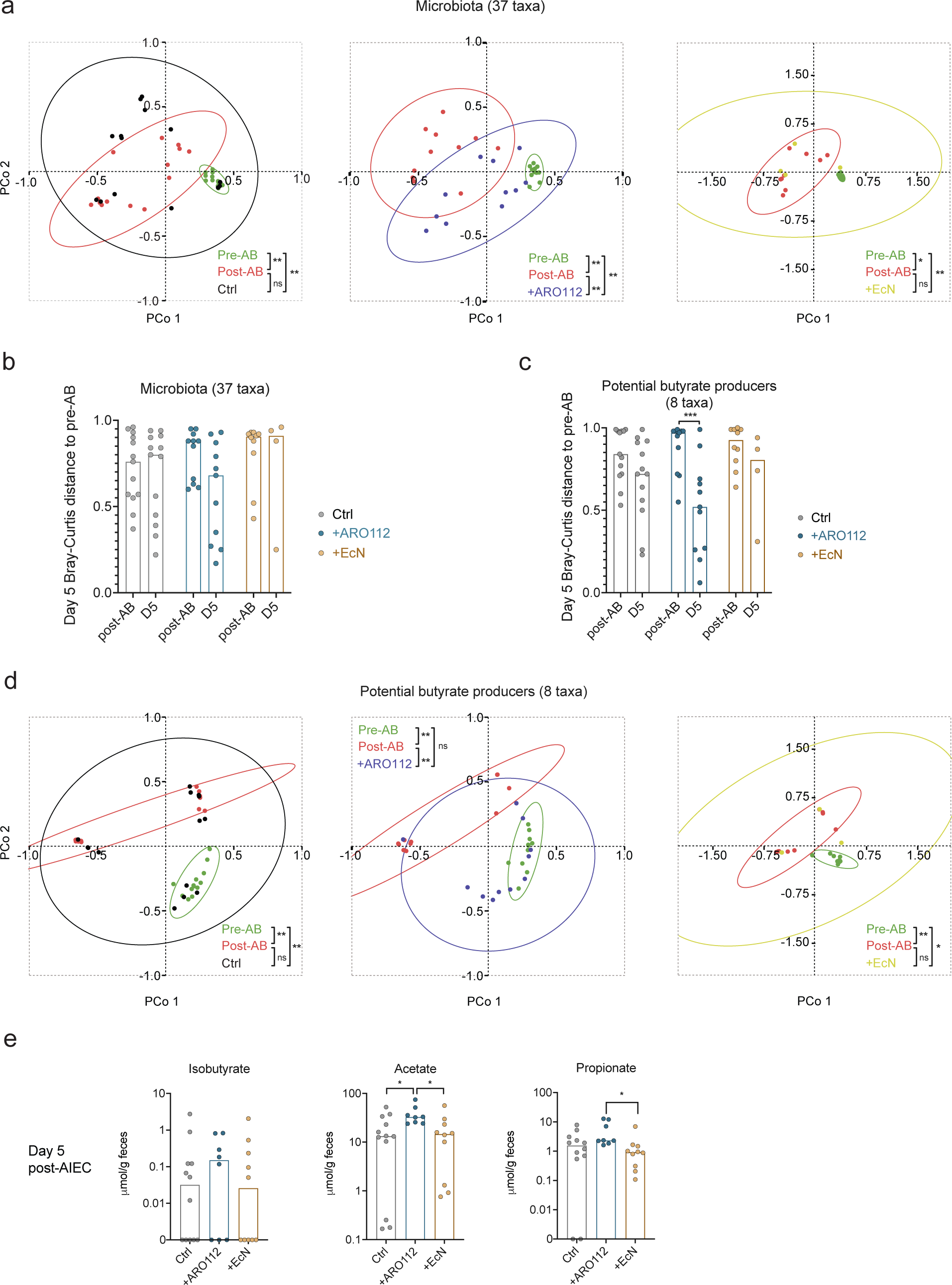
Comparison between treatments of different microbiota-related parameters in experiment in Nod2^-/-^ mice after 4 days of treatment. **a,** Principal Coordinate Analysis of Bray-Curtis dissimilarity index displaying clustering of samples before antibiotic treatment (pre-AB), after antibiotic treatment (post-AB), and after treatment (day 20 of experiment) with PBS (Ctrl; 14 samples) or probiotics +ARO112 (11 samples) or +EcN (4 samples), using microbiota composition (37 taxa). **b,** Analysis of the Bray-Curtis dissimilarity distances, obtained comparing the samples of each mouse pre-AB, to their position post-AB or post-treatment (day 5). **c,** Analysis of the Bray-Curtis dissimilarity distances, obtained comparing the samples of each mouse pre-AB, to their position post-AB or post-treatment (day 5). **d,** Principal Coordinate Analysis of Bray-Curtis dissimilarity index displaying clustering of samples before antibiotic treatment (pre-AB), after antibiotic treatment (post-AB), and after treatment (day 5 of experiment) with PBS or probiotics (+ARO112 or +EcN), for the 8 taxa of potential butyrate producers depleted by antibiotics (*Alistipes*, *Syntrophococcus*, *Ruminococcus*, *Butyricicoccus*, *Dorea*, *Oscillibacter*, unclassified *Ruminococcacea*, unclassified *Lachnospiraceae*). Ellipses in **a** and **d** represent 90% confidence intervals. One-way PERMANOVA with 999 permutations were performed with Bonferroni corrections (**a** and **d**) and two-way ANOVA were tested with Sidak’s correction for multiple comparisons (**b** and **c**) (* p<0.05; ** p<0.01; *** p<0.001).

## Supplementary Information

We analyzed the pathogenic properties that are ubiquitous (present in all 5 strains) or non-ubiquitous (not present in all 5 strains) of each group (Supplementary Table 2). In the non-*pneumoniae Klebsiella* (npK) strains, there are 64 absent properties when compared to *K. pneumoniae* (Kp) strains, indicating an overall lower pathogenic potential predicted for the npK group. The difference in absent predicted pathogenic properties of the npK strains was even higher when compared with *E. coli* (Ec) strains (403), which mainly included genes encoding for transporters, membrane proteins, and genes potentially conferring resistance to antibiotics (Fig. 1b, Supplementary Fig. 1, Supplementary Table 2). There are other properties absent in the npK strains, but present in the Ec strains that are notable, such as genes related to lipopolysaccharide (LPS) biosynthesis and assembly, flagellum, type IV pilus, toxins, ethanolamine utilization, and hemolysin; all of which having been associated with virulence and pathogenesis (Supplementary Table 2)^92–95^.

We investigated the gene products and predicted genome-encoded pathogenic traits in common within each clade, i.e., shared by all 5 strains tested per clade, similarly to the aforementioned analysis for pathogenic properties. A comparison between gene products, which consider the whole genome (Supplementary Table 3), and the predicted genome-encoded pathogenic properties (Supplementary Table 2), shows that the three clades share a higher percentage of predicted pathogenic traits (47.36%) than gene products (14.23%; Extended Data Fig. 1a).

**Supplementary Data Fig. 1.**
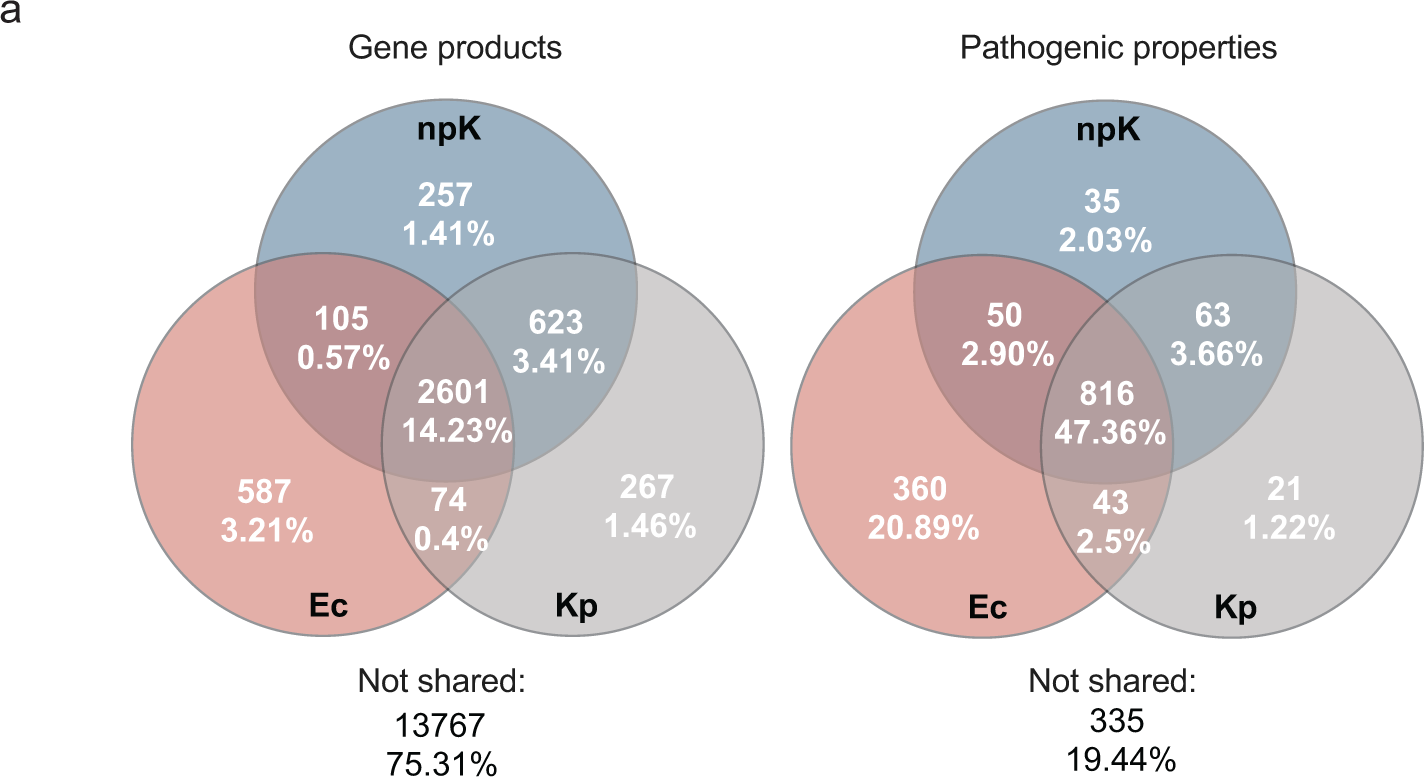
Comparison between gene products and predicted pathogenic properties. Genome analyses were performed in PATRIC BV-BRC software, from where the number of hits for the different virulence factors databases were obtained. **a,** Venn diagrams of total gene products, shared by every member of each clade. **b,** Venn diagram of virulence properties’ hits shared by every member of each clade.

