## Supplementary figures and images for "Gut protective *Klebsiella* species promotes microbiota recovery and pathobiont clearance while preventing inflammation"

### Extended Data Fig.1

Extended Data Fig. 1

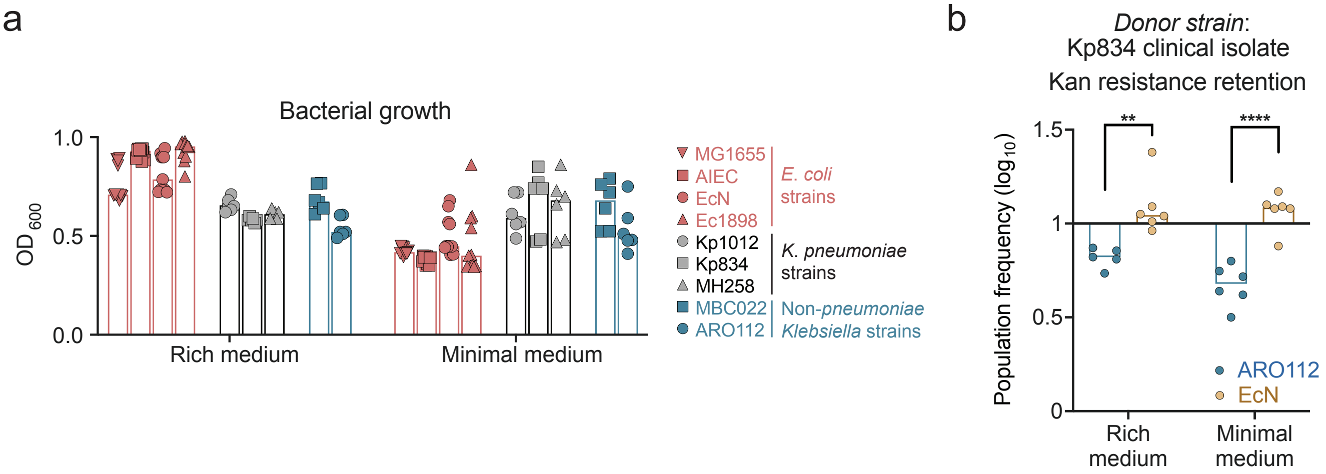

### Extended Data Fig.2

Extended Data Fig. 2

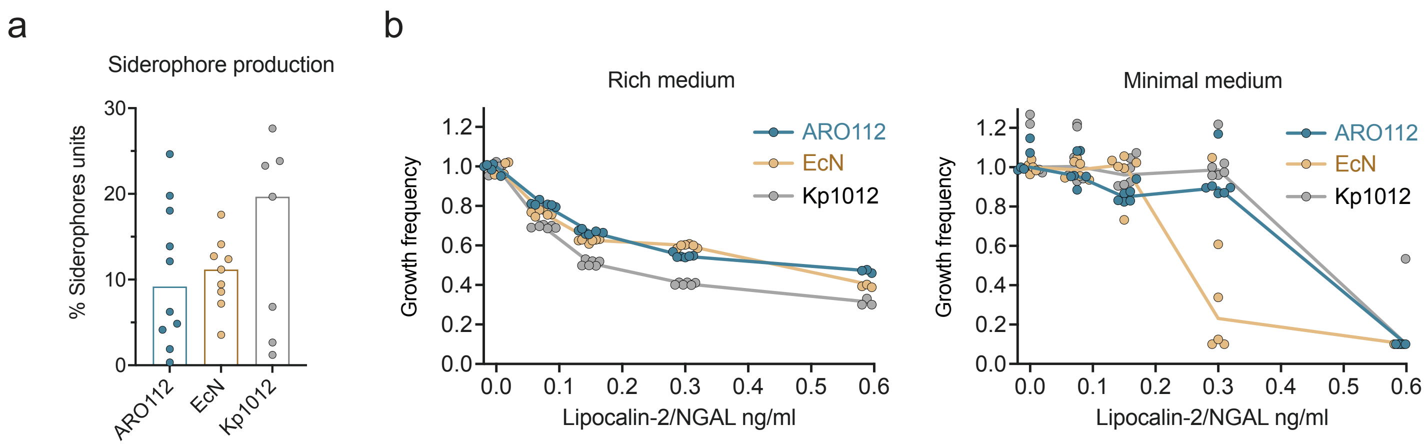

### Extended Data Fig.3

Extended Data Fig. 3

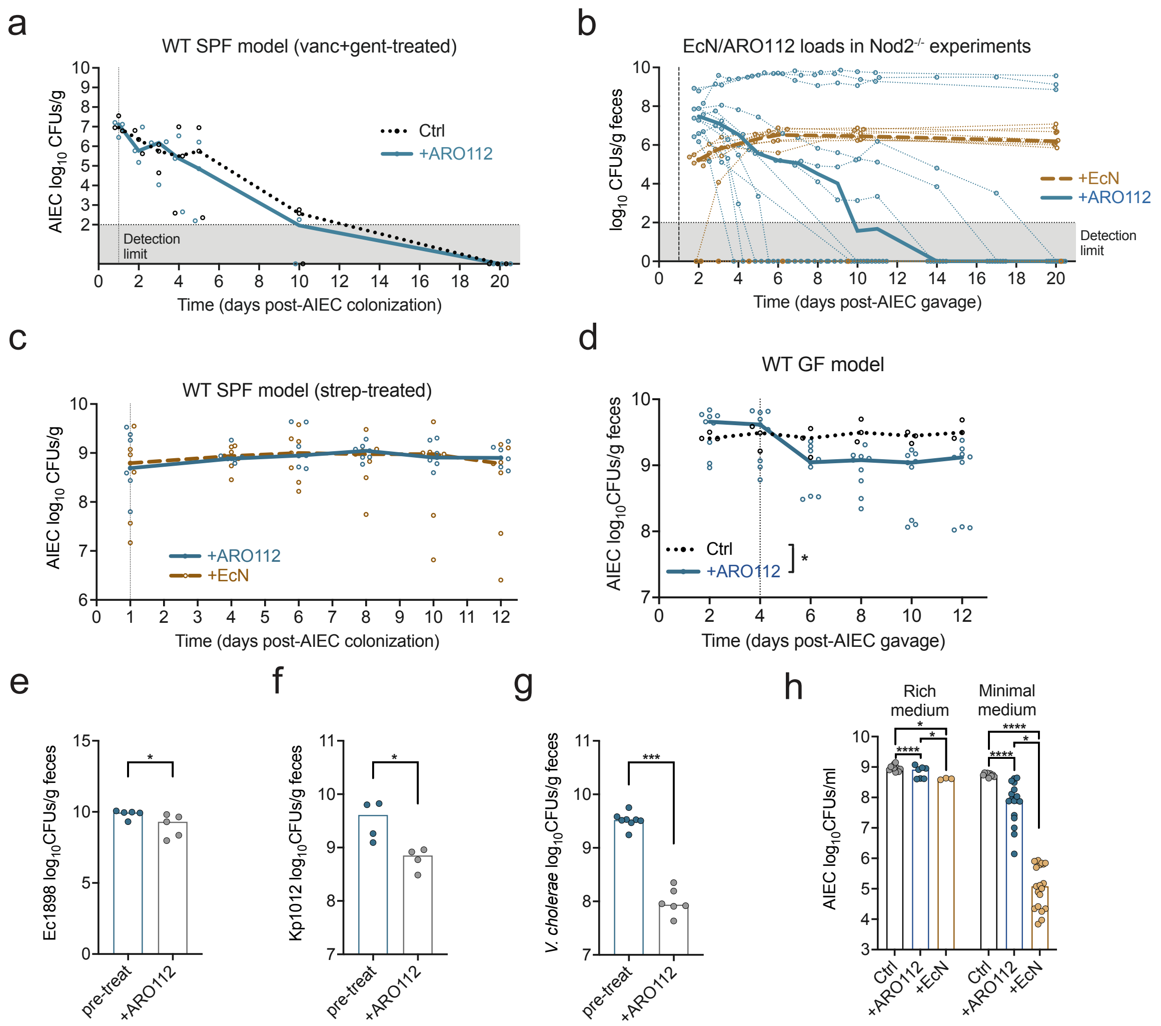

### Extended Data Fig.4

Extended Data Fig. 4

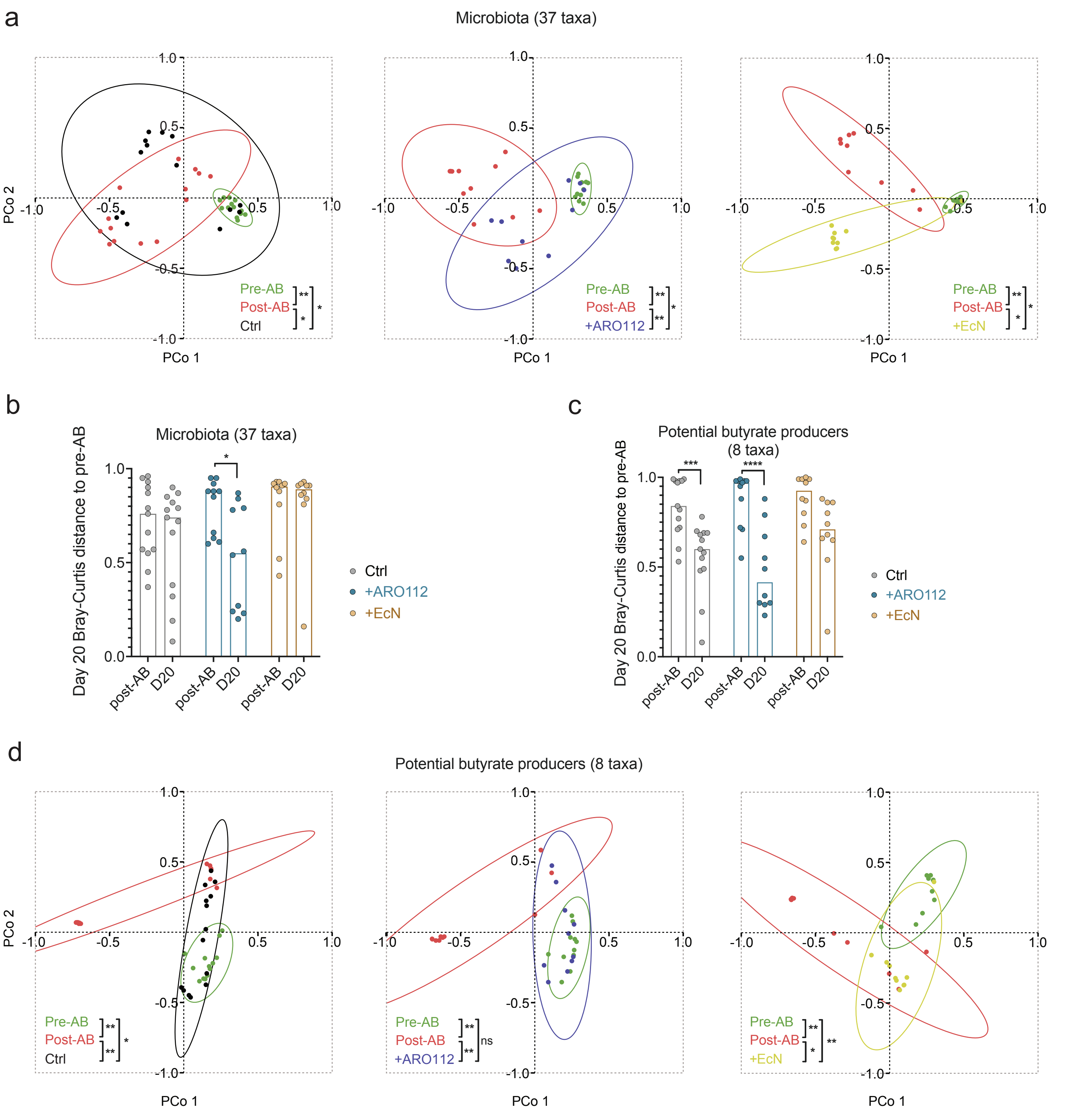

### Extended Data Fig.5

Extended Data Fig. 5

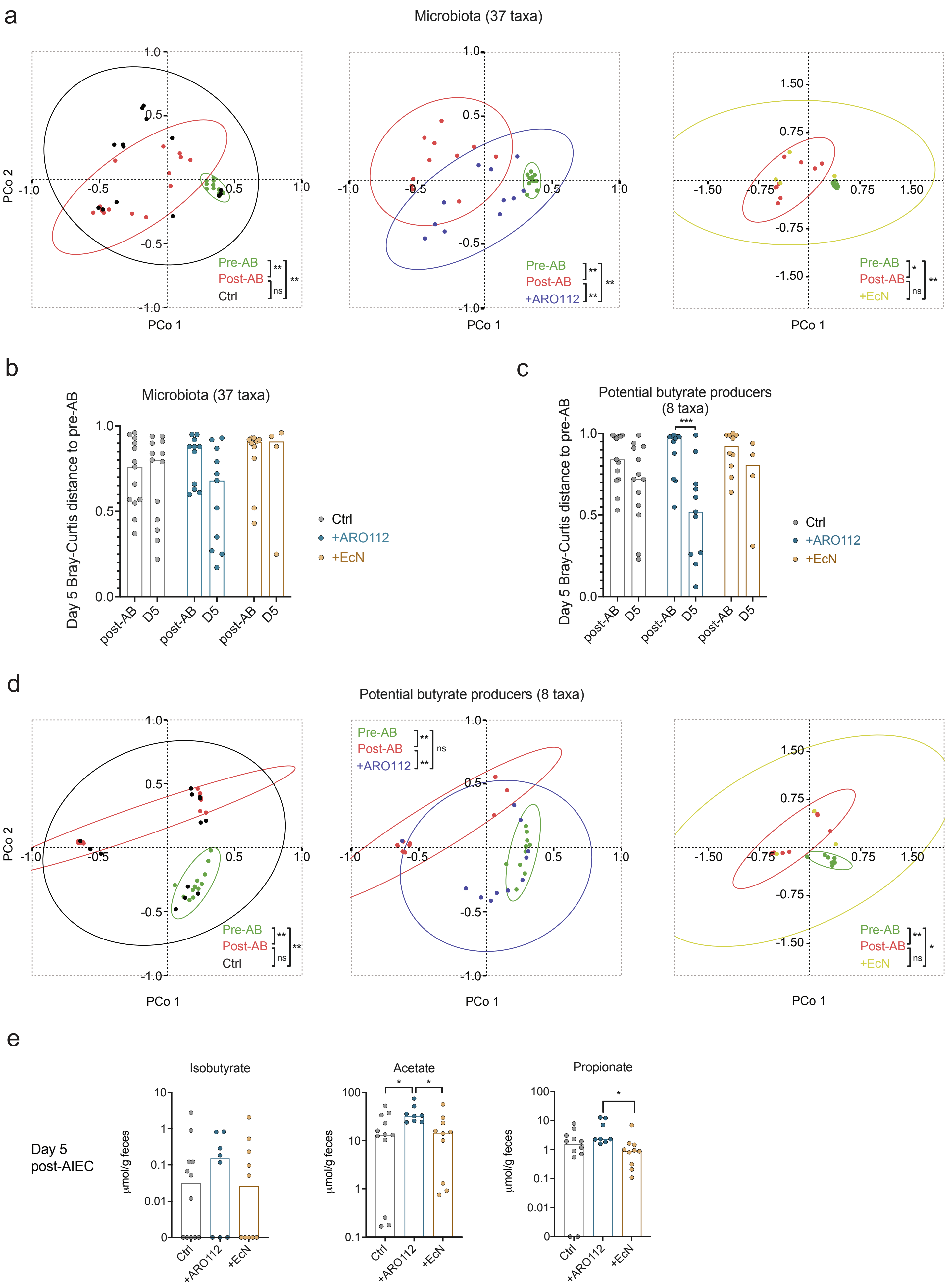

### Supplementary Fig.1

Supplementary Fig. 1

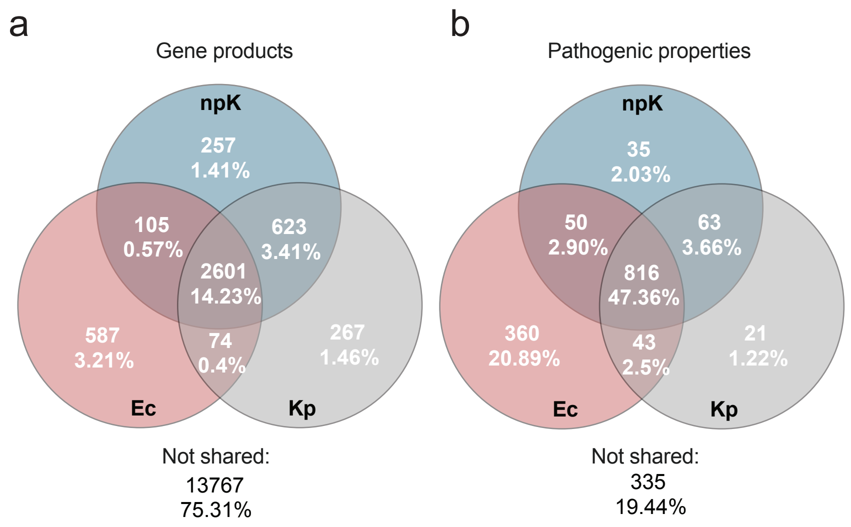
